# Post-GWAS Prioritization of Genome-Phenome Association in Sorghum

**DOI:** 10.1101/2023.12.05.570238

**Authors:** Debasmita Pal, Kevin Schaper, Addie Thompson, Jessica Guo, Pankaj Jaiswal, Curtis Lisle, Laurel Cooper, David LeBauer, Anne Thessen, Arun Ross

**Affiliations:** Department of Computer Science and Engineering, Michigan State University, East Lansing, USA; Translational and Integrated Sciences Lab, Department of Genetics, University of North Carolina Chapel Hill, Chapel Hill, North Carolina, USA; Department of Plant, Soil and Microbial Sciences, Michigan State University, East Lansing, Michigan, USA; Arizona Experiment Station, University of Arizona, Tucson, Arizona, USA; Department of Botany and Plant Pathology, Oregon State University, Corvallis, Oregon, USA; KnowledgeVis, LLC., Altamonte Springs, Florida, USA

**Keywords:** plant phenotype, *Sorghum Bicolor*, maximum canopy height, maximum growth rate, GWAS, SNP prioritization, feature engineering

## Abstract

Genome-Wide Association Studies (GWAS) are widely used to infer the genetic basis of traits in organisms, yet selecting appropriate thresholds for analysis remains a significant challenge. In this study, we developed the Sequential SNP Prioritization Algorithm (SSPA) to elucidate the genetic underpinnings of two key phenotypes in *Sorghum bicolor*: maximum canopy height and maximum growth rate. Utilizing a subset of the Sorghum Bioenergy Association Panel cultivated at the Maricopa Agricultural Center in Arizona, our objective was to employ GWAS with specific permissive-filtered thresholds to identify the genetic markers associated with these traits, allowing for a broader collection of explanatory candidate genes. Following this, our proposed method incorporates a feature engineering approach based on statistical correlation coefficient to reveal patterns between phenotypic similarity and genetic proximity across 274 accessions. This approach helps prioritize Single Nucleotide Polymorphisms (SNPs) likely to be associated with the studied phenotype. Additionally, we evaluated the impact of SSPA by considering all variants (SNPs) as inputs, without any GWAS filtering, as a complementary analysis. Empirical evidence including ontology-based gene function, spatial and temporal expression, and similarity to known homologs, demonstrated that SSPA effectively prioritizes SNPs and genes influencing the phenotype of interest, providing valuable insights for functional genetics research.

## 1. Introduction

Genome-wide association studies (GWAS) [1] are widely used to infer the genetic basis of phenotypic traits in organisms. However, the challenge lies in selecting an appropriate statistically significant threshold (p-value) for the analysis, considering the false discovery rate [2]. A stringent significance threshold for associations could potentially miss causative variance present at lower frequencies in the population. Additionally, a stringent trait correlation filter is likely to be confounded with the population structure. While these features could potentially be highly predictive of the phenotypes, they may not provide insights into underlying biology and gene function, but rather into the breeding history or source of accessions in the population. Therefore, our objective is to utilize certain permissive-filtered GWAS thresholds that retain many more candidate Single Nucleotide Polymorphisms (SNPs) than what the GWAS analysis itself would have identified as significantly associated with the phenotypic trait. This approach would likely increase false positives as well as true positives, leading to subsets of potentially informative SNPs whose association with the phenotypes was presumably not confounded with the population structure.

Building on the genetic markers identified by permissive-filtered GWAS thresholds, we developed the **Sequential SNP Prioritization Algorithm (SSPA)** that employs a feature engineering technique commonly used in Machine Leaning (ML). The core idea is to compare the resemblance of phenotypic similarity − determined by differences in *normalized* phenotypic trait measurements among accessions − with their genetic relatedness using statistical correlation coefficient. This process aids in prioritizing Single Nucleotide Polymorphisms (SNPs) and genes that are likely to influence the phenotype under study. Our algorithm also incorporates an *ordering* of the prioritized SNPs based on their impact on phenotypic traits, addressing the challenge of discovering SNPs with lower effect estimates through traditional GWAS [3]. To evaluate the effectiveness of our method, we focused on the phenotypes of **maximum canopy height** and **maximum growth rate** across 274 accessions of *Sorghum bicolor*, a subset of the Sorghum Bioenergy Association Panel cultivated at the Maricopa Agricultural Center in Arizona. Our code and data are available on GitHub.^1^

We primarily intended to leverage GWAS due to its proven capability in identifying causal SNPs for complex phenotypic traits. Therefore, our method is designed as post-GWAS analysis, following the selection of less stringent GWAS thresholds. In the literature, the most common approach for post-GWAS prioritization of SNPs involves annotating different genomic features, such as Expression Quantitative Trait Loci (eQTLs), sequence-specific DNA-binding factors, with GWAS outputs to identify SNPs for further investigation [4]. This method relies on the physical proximity of a SNP to a known genomic feature to assign relevance; however, while physical proximity can suggest function, it does not necessarily mean that the SNP will affect the phenotype of interest.

Researchers have also proposed other post-GWAS methods to identify most significant loci, which can further be utilized for functional delineation and plant breeding improvement programs [5]. Some approaches were based on computing prioritization scores using p-values, while others utilized meta-analysis, pathway-based analysis, haplotype-based analysis, etc. Cai et al. conducted post-GWAS analysis by combining gene-based association signals and gene expression data to identify putative causal genes for mastitis resistance in dairy cattle [6]. Marina et al. proposed guilt-by-association based prioritization analysis using functional information collected from various sources and exploited fuzzy-based similarity measures to identify functional candidate genes for milk and cheese-making traits in Assaf and Churra dairy breeds [7]. Additionally, ML and deep learning algorithms were also applied for post-GWAS analysis to prioritize disease-associated loci by annotating numerous biological features with GWAS outputs, considering prioritization as a classification problem [8].

In this work, SNP prioritization was achieved using a feature engineering approach that leverages correlation coefficients to compare phenotypic similarity among accessions with their genetic proximity. This contrasts with the p-value-based methods commonly used in existing score-based post-GWAS prioritization schemes. Our goal was to establish associations between genotypes and phenotypes based on statistical correlation rather than regression, which is often used in genome-phenome association studies, e.g., GWAS and any BLUP-based methods [9]. While both regression and correlation are statistical methods for examining relationships between two variables, they can yield different insights depending on the patterns present in the dataset. Regression analysis typically models the relationship between independent and dependent variables to predict the

dependent variable’s value. Conversely, correlation quantifies the strength and direction between two independent distributions, ranging from -1 (perfect negative correlation) to +1 (perfect positive correlation), with 0 indicating no linear relationship. This helps determine if any relationship exists between them at all.

Moreover, instead of analyzing the data at the level of individual accessions, we examined the relationships between every possible pair of accessions. This pairwise approach provides a more detailed understanding of how genetic and phenotypic similarities correspond to each other across accessions, revealing nuanced associations that might be missed when considering accessions individually. Additionally, we utilized *normalized* phenotype measurements to compute the phenotypic similarity among accessions employing a logistic growth curve model, which introduced more resilience to outliers in our experiments and made the extracted results more comparable across different environmental conditions. We also explored the prospect of our proposed algorithm by investigating the differences in results when applying a GWAS filter and when not using any GWAS filter. In summary, the contribution of this work is as follows:

- Developed an algorithm employing a feature engineering technique using statistical correlation for post-GWAS prioritization of SNPs which are likely to affect the phenotype of interest.
- Conducted experiments using 274 accessions of *Sorghum bicolor* grown in Arizona, focusing on maximum canopy height and maximum growth rate phenotypes.
- Assessed the generalizability of our method by utilizing the phenotypic measurements of the same accessions grown in a different environment (South Carolina).
- Investigated the impact of our algorithm in prioritizing SNPs without applying GWAS filter.

## 2. Materials and Methods

### 2.1. Plant Material and Phenotype Dataset Description

The multi-year TERRA Phenotyping Reference Platform (TERRA-REF) program [10,11] studied how the environment and other factors affected the growth of agricultural products. This program aims to maximize growth and, consequently, maximize agricultural bioenergy output. Season 6 of the program was conducted in the summer of 2017, where *Sorghum bicolor* was chosen as the target organism due to its drought tolerance and variety of applications in food, fuel, and fiber, with potential improvement through targeted breeding. A subset of 336 sorghum accessions from the Bioenergy Association Panel (BAP) [12] was cultivated at the Maricopa Agricultural Center (MAC) in Arizona. Throughout the growing season, automated large field scanner systems monitored these plants for phenotype measurements every day or two. In addition, a small number of common phenotypes were also recorded by field scientists.

A suite of measurements, including photographs, canopy height, leaf characteristics, and soil environmental factors, collected regularly by automated systems, were utilized to analyze each accession’s growth patterns [10]. Two replicate plots of each accession were planted within a 200m x 20m field. Data collected by TERRA-REF is available in the public domain, along with experimental design, measurement methods, and other study metadata [11]. Our analysis utilized canopy height measurements which are available for most days during the growing season and whole-genome resequencing data. Plot-level canopy height measurements were estimated from laser scans using an algorithm calibrated to hand measurements.

### 2.2. Phenotype Data Preparation and Normalization

Automated measurements of canopy height over the growing season were fitted with logistic growth curves to obtain the phenotypic traits of *maximum canopy height* and *maximum growth rate* relative to growing degree days (gdd), defined as the accumulation of heat energy above 10°C following the “Method 1” described by McMaster & Wilhelm [13]. This normalization approach can yield phenotypic traits that are less sensitive to outliers, more comparable across environmental conditions, and permit greater biological inference.

First, we applied a cleaning algorithm that excluded accessions with fewer than 35 automatic measurements per season or measurements on fewer than 40% of days, resulting in a total of 326 accessions. This preprocessing step yielded a time series that could robustly estimate the parameters of two phenotypes.

Time series of canopy height and growing degree days (gdd) for each accession were modeled in a Bayesian framework with the likelihood:

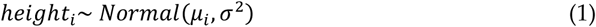

where, *μ* is the expected value of height, *σ*^2^ is the measurement error variance, and *i* indexes each observation. The logistic model was defined as:

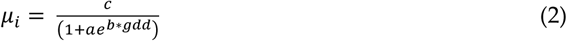

with the logistic parameters a, b, and c and the covariate gdd associated with each observation. However, we re-parameterized the model to obtain more biologically meaningful parameters by defining:

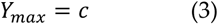

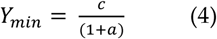

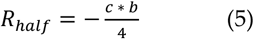

Finally, we transformed the logistic parameters to the whole real line, which allowed for faster mixing and improved model convergence:

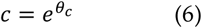

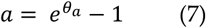

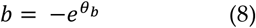

All root nodes were given wide, relatively non-informative standard priors, including *Normal(0,1000)* for all transformed logistic parameters (θ_a_, θ_b_, θ_c_) and Gamma(0.1, 0.1) for the observation precision 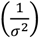, where Normal() and Gamma() refer to probability distributions.

We implemented the above model in JAGS 4.3.0 [14] via R/rjags [15,16]. Three parallel Markov Chain Monte Carlo (MCMC) sequences were assigned with dispersed starting values. The models were run until convergence was achieved at a Gelman and Rubin diagnostic parameter of < 1.2 [17]. An additional run of 10,000 iterations with the thinning parameter as 10 yielded a total of 3,000 relatively independent posterior samples for each parameter of interest. *The posterior median of Y_max_ (maximum canopy height, cm) and R_half_ (maximum growth rate, cm/gdd) of the accessions were considered as phenotype measurements in our proposed method.* Figure 1 shows the logistic growth curves obtained for two sample accessions. The data and code for the logistic growth curve model are publicly available [18].

**Figure 1.**
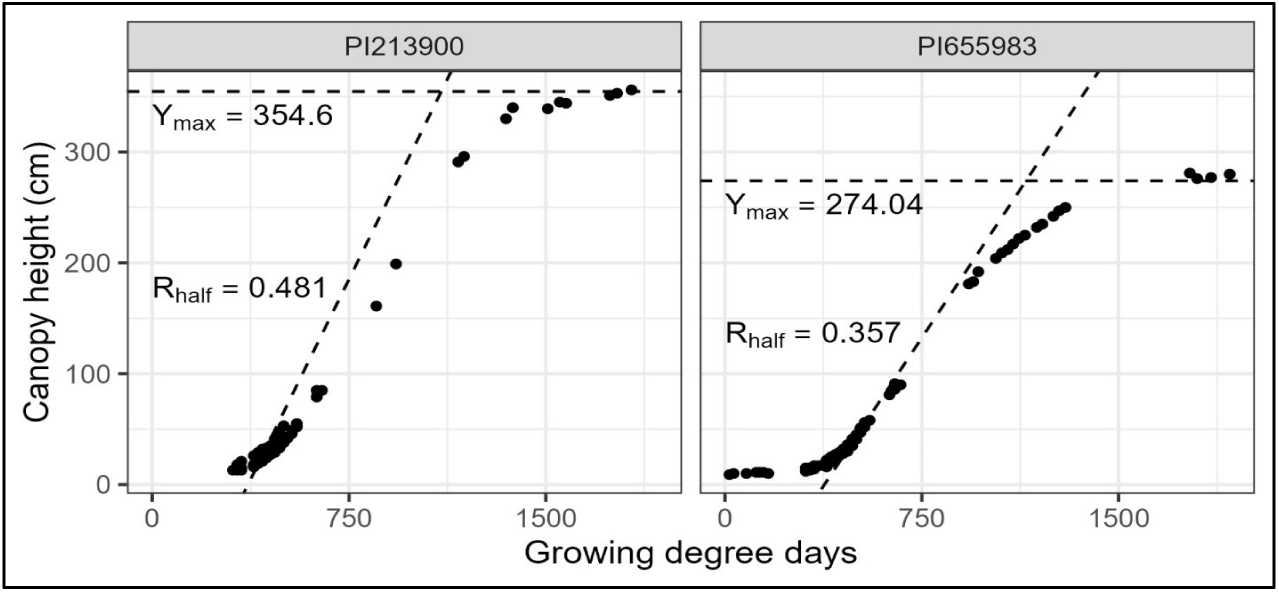
Examples of logistic growth curves to illustrate phenotypes of two sorghum accessions. Using automated canopy heights for each accession, logistic growth curves were fitted and produced maximum canopy height (Y_max_) and maximum growth rate (R_half_), which are indicated by two dashed lines. The cleaning algorithm ensured sufficient observations to produce confident estimations of the parameters.

### 2.3. Computation of Phenotype Similarity Matrix

Based on the *normalized* phenotypic measurements of the sorghum accessions, we computed the *phenotype similarity matrix* separately for each phenotype: maximum canopy height and maximum growth rate. We assumed that the accessions with close genetic proximity would exhibit smaller differences in their phenotypic observations. Here, each phenotype similarity matrix represents the potential correlation (similarity) among accessions. The rows and columns of these matrices correspond to the accessions, and the values indicate the similarity between pairs of accessions, ranging from 0 to 1. A higher similarity value indicates greater similarity between the accessions based on the phenotype under study. The similarity value (*s*_ij_) between the accessions *i* and *j* with phenotype measurements *p_i_*and *p_j_*, respectively, was calculated as follows:

- *Step 1:* The difference in their phenotypic measurements was computed.

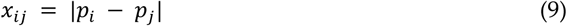

- *Step 2:* Min-max scaling was applied to the difference (x_ij_), using the following equation, to scale the difference in the range [0,1]:

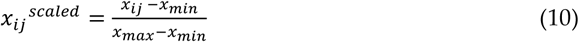

where, *x*;_min_ and *x*;_max_, respectively, denote the minimum and maximum values obtained over all x_ij_’s.

- *Step 3:* To obtain similarity value, the scaled difference was subtracted from 1, so that a higher value indicates a greater similarity between accessions.

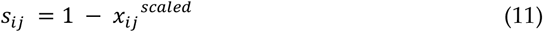

2.4. *Genomic Data Preparation and GWAS:*

We utilized the Sorghum BTX623 reference genome v3.1 for our study. Whole genome sequencing data was obtained from the TERRA-REF project [11], where 384 sorghum accessions from the sorghum Bioenergy Association Panel (BAP) were sequenced with ∼25x coverage. Shotgun sequencing (127-bp paired-end) was conducted using an Illumina X10 instrument at the HudsonAlpha Institute for Biotechnology. Following the alignment of sequence reads to the BTX623 reference genome v3.1 (available from Phytozome [19]), we downloaded GVCF files with variant calls from CyVerse, merged them, and applied filtering protocols as outlined in the TERRA-REF documentation. SNPs were filtered using GATK Variant Filtration, and accessions lacking phenotypic observations were removed. Only biallelic sites were retained. *This set of SNPs was utilized while exploring the impact of our proposed algorithm SSPA without GWAS filtering*.

Next, GWAS was conducted using rMVP [20] with the FarmCPU algorithm [21] for each normalized phenotype (maximum canopy height and maximum growth rate), incorporating a kinship matrix and one principal component to account for relatedness and population structure. The results were filtered with p-values of .0001, .0005, and .001, resulting in three distinct lists of SNPs. Additionally, these lists were cross-referenced with QTL regions known to be associated with height and growth obtained from the Sorghum QTL Atlas [22]. *This process yielded six SNP lists for each phenotype, which were used to filter the GVCF file, resulting in a total of 12 VCF files corresponding to the two phenotypes, hereafter, referred to as experimental conditions.* Detailed data and links to download the VCF files, both before and after GWAS, are available in the Supplementary File.

### 2.5. Creation of SNP Similarity Matrices

Subsequently, each VCF file corresponding to an experimental condition, was processed to create SNP similarity matrices, comparing the allelic similarity of every pair of accessions at that chromosomal location, hence each SNP similarity matrix corresponds to a single SNP. The values of these similarity matrices were computed with an addition of 0.5 for each matching allele between any pair of accessions, while reversed heterozygous SNPs received a value of 1.0. For instance, if a VCF file consists of ‘N’ SNPs across ‘a’ number of accessions, there would be ‘N’ SNP similarity matrices. Each of these similarity matrices is a square matrix of dimensions ‘a x a’, with values encoded as one of three discrete values: {0, 0.5, 1.0}.

From the 326 accessions obtained as part of the logistic growth model, we used 274 accessions that were common to both Seasons 4 and 6 of the TERRA-REF program for generating the similarity matrices.^2^ Thus, each SNP similarity matrix in our experiment has dimensions of 274 x 274.

### 2.6. Sequential SNP Priotization Algorithm

Thereafter, we aimed to obtain a set of prioritized SNPs *individually for each experimental condition* by comparing the corresponding *SNP similarity matrices* with the respective *phenotype similarity matrix*. The objective was to identify SNP similarity matrices which closely resemble the phenotype similarity matrix, i.e., SNPs wherein the genetic variations between the accessions aligns with the variations in the phenotypic measurements. To achieve this, we developed **Sequential SNP Prioritization Algorithm (SSPA**) using a feature engineering approach. This algorithm is inspired by the concept of Sequential Forward Selection (SFS) [23]. We employed the Pearson correlation coefficient (*ρ*_(X,Y)_) to measure the similarity (relationship) between the two data matrices. It is defined by the covariance of two data matrices X and Y, divided by their standard deviations.

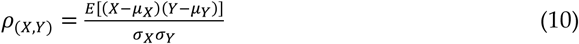

where, *μ*_X_ and *μ*_Y_ are, respectively, the mean and, *σ*_X_ and *σ*_Y_ are, respectively, the standard deviations of the data matrices X and Y.

SFS is a dimensionality reduction technique commonly used in ML to manage high-dimensional datasets. Its purpose is to extract a subset of features that contribute most to the prediction performance of the model. The method operates *iteratively*, selecting the most important features one-by-one based on an evaluation criterion and appending them to an initially empty candidate list. The stopping criteria can be defined as a pre-set number of features (i.e., the desired number of selected features) or the point at which the addition of further features no longer improves the evaluation criterion.

Our proposed method SSPA works in a similar fashion: We initiated the process by defining a “*maximum desired number of prioritized SNPs*” (e.g., 1000), and an empty candidate list. In the first iteration, we selected a SNP similarity matrix from the set of SNP similarity matrices that resulted in the highest correlation coefficient with the phenotype similarity matrix and added it to the candidate list. Starting from the second iteration, we selected the SNP similarity matrix from the remaining set (those not present in the candidate list) that, when summed up with all the similarity matrices together (matrix summation) in the candidate list produced the highest correlation coefficient with phenotype similarity matrix. This selected matrix was then appended to the candidate list. This process was continued iteratively, each time seeking the *next most cumulatively impactful* SNP similarity matrix and adding it to the candidate list until the number of matrices in the candidate list reached the “maximum desired number of prioritized SNPs”. During the execution, we also recorded the highest correlation coefficient obtained at each iteration in a list, allowing us to determine the iteration index (K) with the greatest correlation coefficient. Based on K, we obtained a set of prioritized SNPs by selecting the first ‘K’ SNP similarity matrices from the candidate list. Therefore, at the end, our method generates a *final SNP similarity matrix* (sum of ‘K’ number of prioritized SNP similarity matrices) based on the allelic proximities among accessions maximizing the correlation coefficient with the phenotype similarity matrix. The algorithm is described below:

#### Input

- *‘N’ no. of SNP similarity matrices (each SNP corresponds to a similarity matrix) involving ‘a’ unique accessions: S_j_* ∈ *{0.0, 0.5, 1.0}, where j =1, 2,…,N; each S_j_ is of dimension a x a*.
- *Phenotype similarity matrix ‘P’ across the same ‘a’ number of accessions with dimension a x a*
- *Maximum desired number of prioritized SNPs: n*

#### Output

- *‘K’ no. of prioritized SNPs and K<=n*

***Initialize:*** *Candidate list: L =*⊘ *; Correlation coefficient list: C =*⊘

#### Algorithm

- ***Step 1:*** *Compute the Pearson correlation coefficient of each S_j_ with P, for j = 1, 2,…,N, and select Sj* with highest correlation coefficient*.
- ***Step 2:*** *Append Sj* to L and highest correlation coefficient value to C*.
- ***Step 3:*** *Compute the matrix summation of all the similarity matrices in L and Sj, and calculate the Pearson correlation coefficient with P, where j* ∈ *{1,2,…,N} and S_j_* ⊄ *L*
- ***Step 4:*** *Select S_j_ from Step 3 that resulted in the highest correlation coefficient value and append it to L and highest correlation coefficient value to C*.
- ***Step 5:*** *Repeat Steps 3 and 4 until the number of SNP similarity matrices in L reaches the maximum desired number of prioritized SNPs n*.
- ***Step 6:*** *Select the index K of the list C with maximum correlation coefficient*.
- ***Step 7:*** *Select the first K number of SNP similarity matrices from L (i.e., the summation of which, termed as final similarity matrix, resulted in the maximum correlation coefficient value with P). These SNP similarity matrices correspond to K prioritized SNPs*.

Figure 2 presents a streamlined flowchart of this algorithm. This process enables the cumulative identification of prioritized SNPs through a sequential evaluation of both local and global relationships with phenotypic trait, ultimately ranking them by their influence on the studied phenotype. In the functional analysis of these prioritized SNP sets, we refer to the term “Top k” to indicate the first ‘k’ SNPs in the prioritized SNP list.

**Figure 2.**
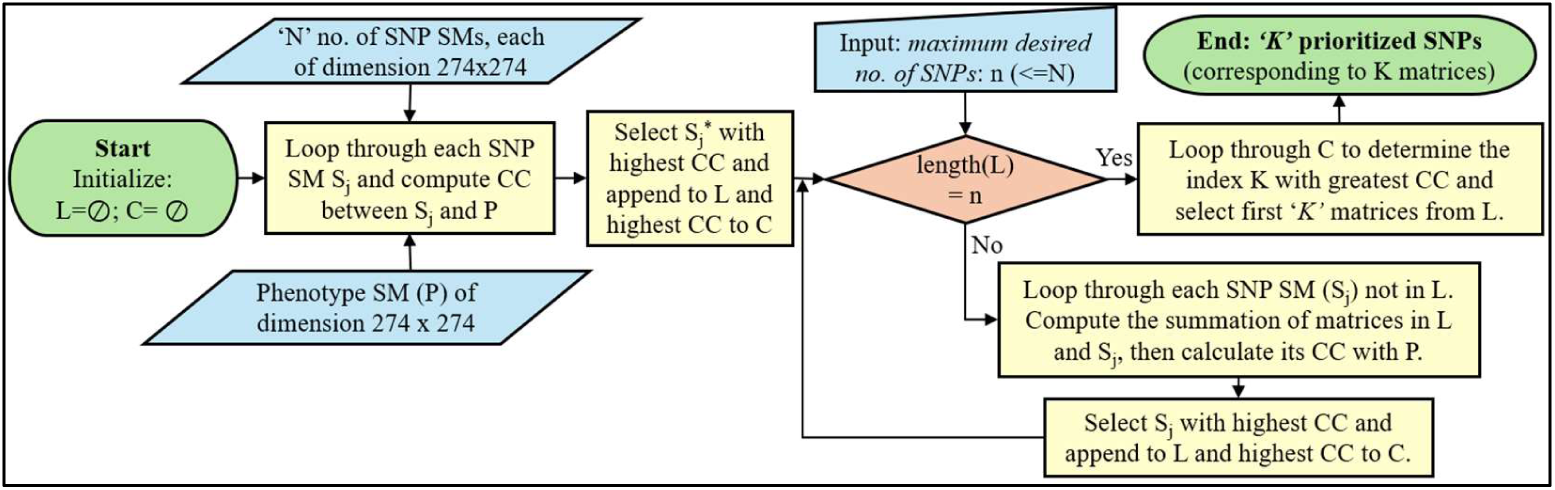
Flowchart of Sequential SNP Prioritization Algorithm (SSPA). This algorithm inputs a single phenotype similarity matrix and multiple SNP similarity matrices to determine a set of prioritized SNPs (L: Candidate List, C: Correlation Coefficient List, SS: Similarity Matrix, CC: Correlation Coefficient).

### 2.7. Prioritizing SNPs without GWAS filtering

As a complementary analysis, we explored the impact of our proposed algorithm SSPA when applied to all variants (SNPs) without applying any GWAS filtering. For this experiment, we only focused on the maximum canopy height phenotype. Due to the large volume of data and limited computational resources, we were unable to execute our method on the full GVCF file at once. Therefore, we created 10 VCF files, each containing SNPs corresponding to one of the chromosomes (detailed data available in the Supplementary File). The goal was to first process each VCF file individually to prioritize SNPs using SSPA following the creation of SNP similarity matrices from each chromosome. Next, we merged the prioritized lists of SNPs from each chromosome together and executed the SSPA on the SNP similarity matrices that correspond to the combined set of prioritized SNPs. This enabled us to obtain a final set of prioritized SNPs without GWAS filtering for the analysis.

### 2.8. SNP Annotation and Function

The prioritized SNPs extracted by SSPA were annotated with overlapping or nearest known gene identifiers using the SnpEff package [24]. The analysis of function was carried out on either Phytozome [19], Gramene [25], Planteome [26] or the Gene Ontology [27] online databases. A list of unique gene identifiers was submitted for Gene Ontology (GO) Enrichment Analysis using PANTHER 17.0 [28] and the Sorghum bicolor reference list [29]. We performed separate enrichment analyses using the complete set of GO Biological Process, Molecular Function and Cellular Component annotation datasets. Fisher’s exact test with FDR correction was employed for the analyses.

The genes identified that neighbor the “Top 10” SNPs (from SnpEff) for both phenotypes with the GWAS filtering (with and without QTL filtering) at p=.0001 were further explored for their functional role by using Gramene and the EMBL-EBI Gene Expression Atlas. This was also done for the “Top 10” SNPs for the height phenotype obtained without the GWAS filtering. InterPro protein domain annotations [30] were obtained by querying Gramene BioMart [31,25]. The gene IDs were also used to query the EMBL-EBI Gene Expression Atlas [32] to explore their baseline gene expression. The baseline expression data of genes showing a default minimum expression level of 0.1 transcripts per million (TPM) were downloaded for experiments E-MTAB-3839 [33], E-MTAB-4400 [34], and EGEOD-98817 [35]. Expression heatmaps were generated using ClustVis (Beta) [36]. The Sorghum QTL data was queried from the Sorghum QTL Atlas [22] and mapped on Gramene’s sorghum genome browser. The SNPs were also queried using Gramene’s variant effect prediction (VEP) tool [37,38] to find any overlapping genes or existing SNPs in that database.

### 2.9. Generalizability Testing

To validate the generalizability of our proposed method in a different environment, we considered *end-of-season height data* of the same accessions grown in Clemson, Florence County, South Carolina [12]. The goal was to examine the associations (relationships) of already extracted prioritized SNPs with the phenotypic observations obtained in a new environment.

We created a *phenotype similarity matrix* using the end-of season phenotypic measurements from Clemson following the process described in Section 2.3.^3^ Next, we calculated the Pearson correlation coefficient between this phenotype similarity matrix and the *final SNP similarity matrix* (i.e., the summation of similarity matrices corresponding to the prioritized SNPs extracted), obtained for each experimental condition based on the maximum canopy height of the MAC Season 6 dataset. A similar validation was also conducted with the final set of prioritized SNPs extracted without GWAS filtering.

## 3. Results

### 3.1. Experiments on post-GWAS Prioritization of SNPs

Figure 3 outlines our proposed approach for post-GWAS prioritization of SNPs. For each experimental condition that corresponds to a specific phenotype and GWAS filtering threshold (described in Section 2.4), we obtained a set of prioritized SNPs using the SSPA. Table 1 summarizes the number of GWAS-filtered SNPs used as input to our algorithm and the respective results. In cases where the total number of GWAS-filtered SNPs (input) exceeded 1000, the *maximum desired number of prioritized SNPs (n)* was set to 1000 during the execution of SSPA. We observed that this limit did not impact the prioritization of SNPs. The highest correlation coefficient i.e., the correction coefficient between phenotype similarity matrix and the final SNP similarity matrix (summation of SNP similarity matrices corresponding to prioritized SNPs) was consistently achieved at an index below 1000 for all experimental conditions. Figure 4 illustrates the trend of the correlation coefficient over the iterations while adding SNP similarity matrices one-by-one to the candidate list for the experiment based on maximum canopy height with a p-value of .001 without QTL filtering.

**Figure 3.**
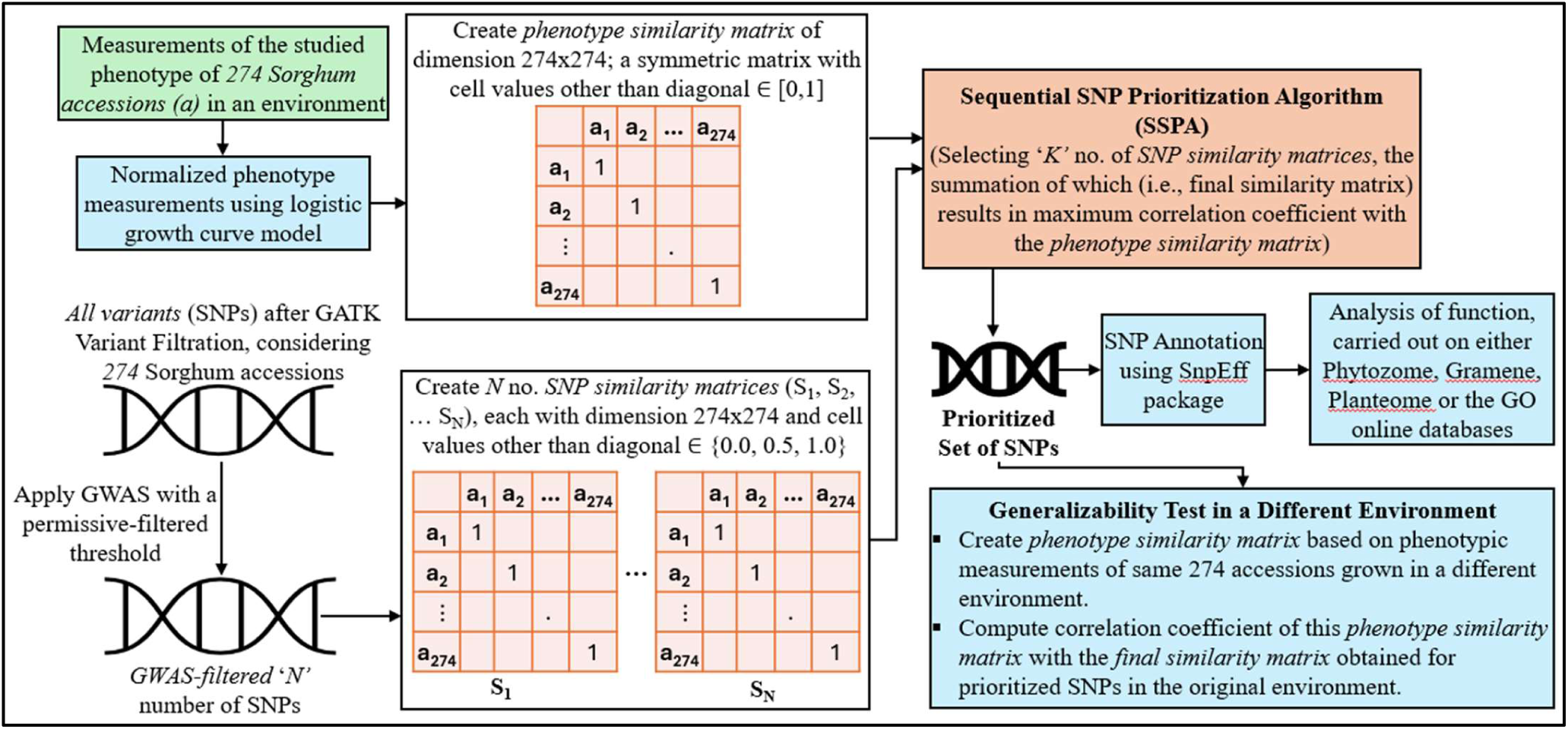
Method pipeline for post-GWAS prioritization of SNPs and its analysis. This diagram illustrates the process for a single experimental condition, beginning with the measurements of the studied phenotype for a set of accessions and GWAS-filtered SNPs. Our proposed method resulted in a set of prioritized SNPs, which were further utilized for functional analysis and to test the generalization capability of our proposed algorithm SSPA.

**Figure 4.**
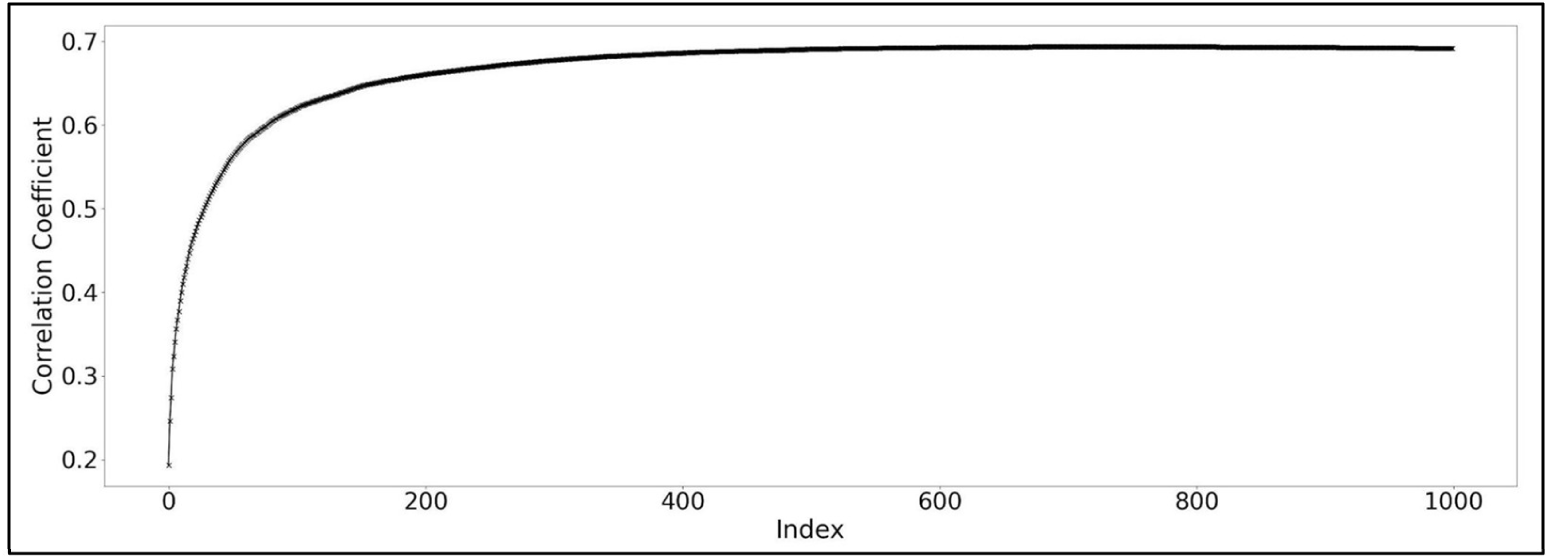
Trend of correlation coefficient during the sequential addition of SNP similarity matrices to the candidate list over the iterations (index) in the execution of SSPA, recorded in Correlation Coefficient List. This analysis pertains to the phenotype of maximum canopy height with GWAS filtering applied at a p-value of .001. As the *maximum desired number of SNPs* was set as 1000 in our experiment, SSPA aimed to select a maximum of 1000 SNP similarity matrix to the candidate list iteratively. The highest correlation coefficient was observed at index 755, resulting into 755 prioritized SNPs for this experimental condition.

**Table 1.**
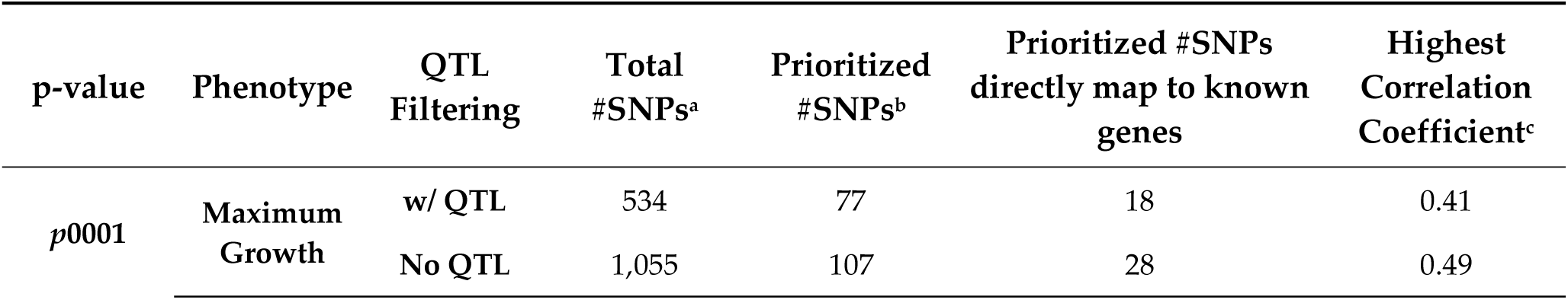
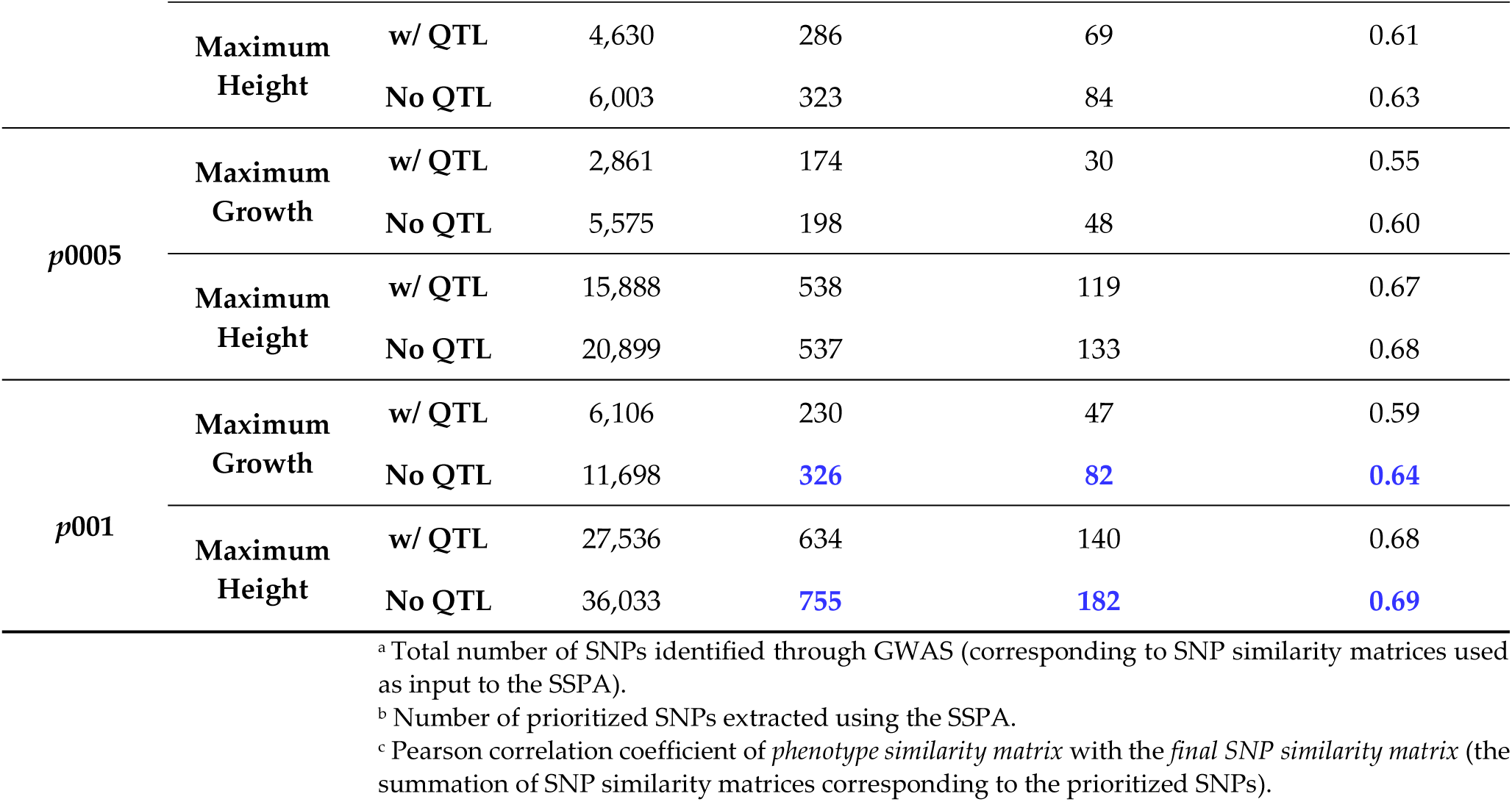
Experimental summary of post-GWAS prioritization of SNPs using permissive-filtered GWAS thresholds.

### 3.2. Experiment on SNPs Prioritization without GWAS filtering

As described in Section 2.7, we first performed chromosome-wise execution of the SSPA on all SNPs without GWAS filtering, setting the *maximum desired number of prioritized SNPs (n)* to 1000. After merging the extracted prioritized SNPs from each chromosome, we executed the SSPA on this combined set to obtain a final set of prioritized SNPs for the analysis. While applying the algorithm to the combined prioritized set, we iterated through the entire list as the total number of SNPs was already reduced during chromosome-wise execution. In this case, the *maximum desired number of prioritized SNPs (n)* was set to the total number of SNPs in the set. Figure 5 shows the trend of correlation coefficient achieved during the addition of SNP similarity matrix to the candidate list while iterating through the combined prioritized set of SNPs obtained from all chromosomes.

**Figure 5.**
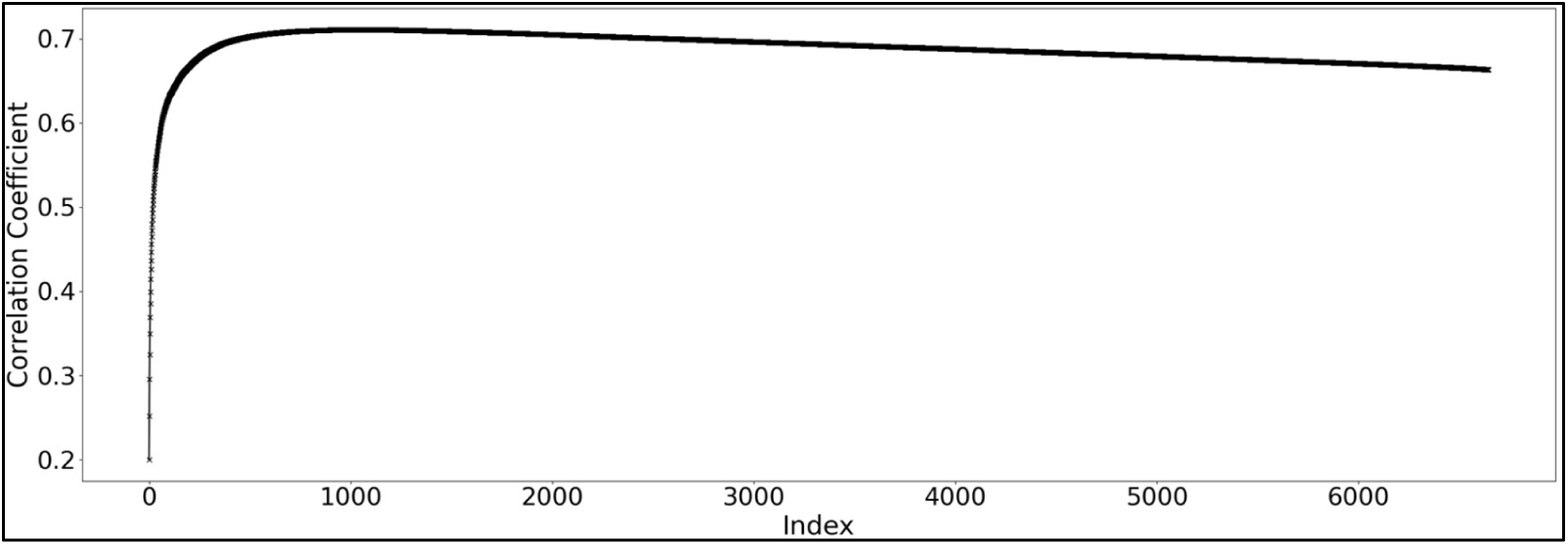
Trend of correlation coefficient during the sequential addition of SNP similarity matrices to the candidate list over the iterations (index) in the execution of the SSPA, recorded in Correlation Coefficient List. This analysis pertains to the combined set of prioritized SNPs corresponding to phenotype of maximum canopy height without GWAS filtering. In this case, the *maximum desired number of SNPs* was set as the total number of input SNP similarity matrices, causing SSPA to iterate through the entire list. The highest correlation coefficient was observed at index 987, resulting into 987 prioritized SNPs for this experiment.

### 3.3. Observations on Prioritized SNPs obtained from Different Experiments

For both phenotypes, the SSPA resulted in fewer prioritized SNPs with a smaller *p*- value, as expected, when applied with GWAS filtering (Table 1 - Prioritized #SNPs). Additionally, filtering by QTLs reduced the number of prioritized SNPs; however, it did not improve the highest correlation coefficient achieved (Table 1 - Highest Correlation Coefficient). *The highest correlation coefficient was achieved at p=0.001 without QTL filtering for both the phenotypes.* Figure 6 describes overlap between prioritized SNPs retrieved using SSPA for different experimental conditions with GWAS filtering and without GWAS filtering using UpSet plot [39]. We observed the following:

a) A minimal overlap in prioritized SNPs exists between the two phenotypes of height and growth, and this overlap occurred only at the two less stringent p-values of 0.0005 and 0.001 (Figure 6B,6C). At p=0.0001, there was no overlap between the prioritized SNPs for height and growth.
b) Lists of prioritized SNPs with and without QTL filtering have more overlap for the height phenotype, whereas the lists of prioritized SNPs with and without QTL filtering have a moderate amount of overlap for the growth phenotype (Figure 6A, 6B, 6C).
c) At each p-value and for each phenotype, the QTL filtering added between 18% and 31% more unique prioritized SNPs.
d) Most of the prioritized SNPs extracted without GWAS filtering were not identified as informative compared to results obtained with GWAS filtering for maximum height (Figure 6-D).

**Figure 6.**
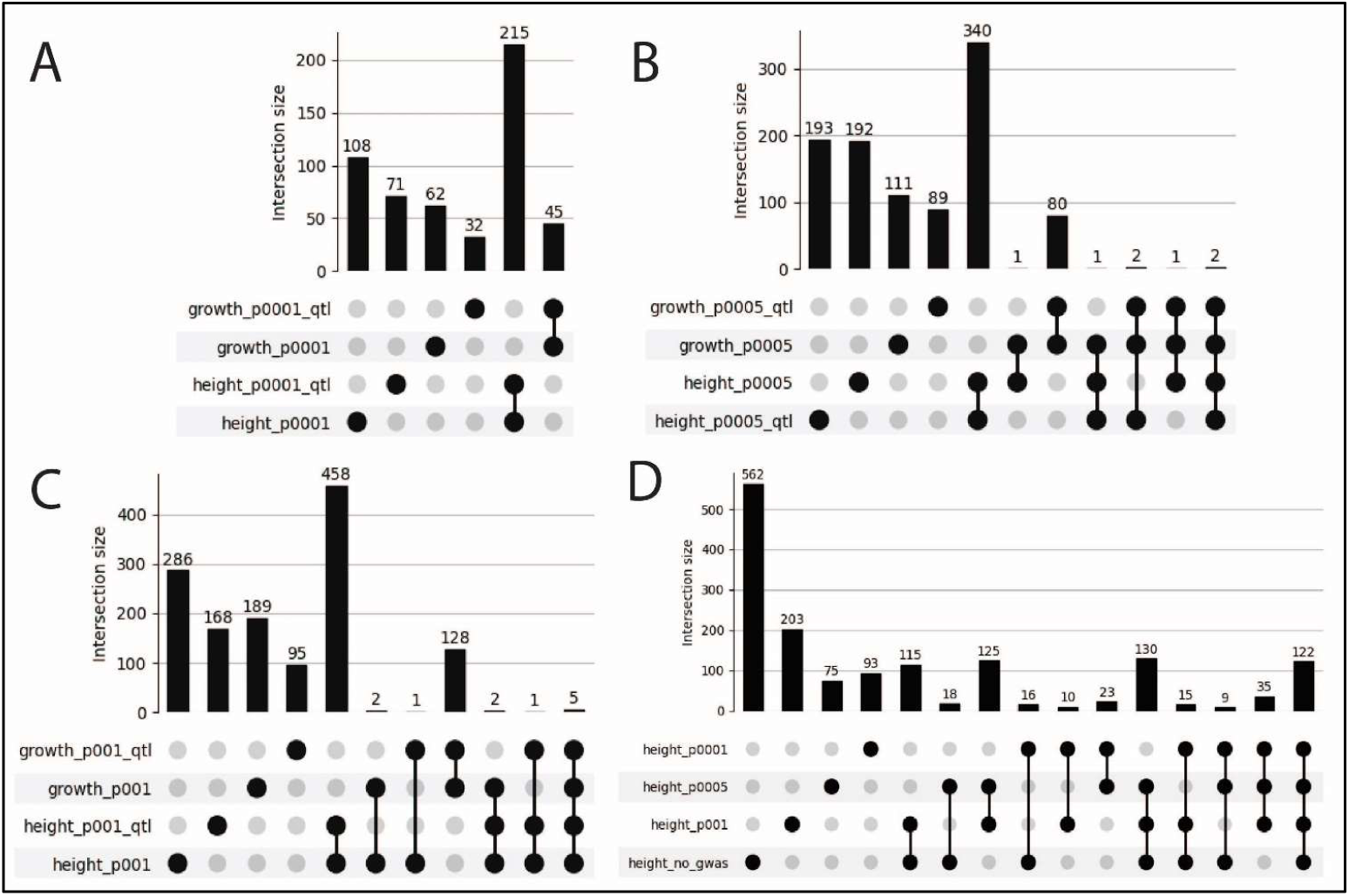
UpSet plot showing the overlap of the prioritized SNPs extracted using SSPA for different experimental conditions with GWAS filtering and without GWAS filtering. It helps visualizing the intersections between the sets in a matrix, where each row corresponds to a set (SNP list) and each column corresponds to a possible intersection with the vertical bar showing the size of the intersection. The filled cells in each column show which set is part of the intersection. Figure was created with UpSetPlot v0.8.0 (https://github.com/jnothman/UpSetPlot). (A) Prioritized SNPs summary for height and growth phenotypes with GWAS filtering at p0.0001. No overlap in prioritized SNPs exists between the phenotypes. (B) Prioritized SNPs summary for height and growth phenotypes with GWAS filtering at p0.0005. There is a total of 7 SNPs in common between the phenotypes. (C) Prioritized SNPs summary for height and growth phenotypes with GWAS filtering at p0.001. There are 11 SNPs in common between the phenotypes. (D) Prioritized SNPs summary for height phenotype with and without GWAS filtering. Most of the SNPs identified as prioritized without the GWAS filtering had not been identified when using the GWAS filtering.

### 3.4. Analysis of Function of Prioritized SNPs

Few prioritized SNPs were mapped to known genes (Table 1 and Table 2), with the majority mappings to intergenic regions. Enrichment analysis showed that 43 unique GO terms related to biosynthetic processes, metabolic processes, and gene expression were over-represented in the annotations of the genes associated with the prioritized SNPs. The triterpenoid biosynthetic process (GO:0016104) and triterpenoid metabolic process (GO:0006722) were the most over-enriched (20.13-fold enrichment) across all the prioritized SNP lists for height phenotype and were in the prioritized SNP lists for the growth phenotype with GWAS filtering at p=0.001 with and without QTL filtering. Nitrogen compound metabolic process (GO:0006807) was one of the few under-enriched (0.72-fold enrichment) GO annotations and was only present in prioritized SNP lists for the height phenotype at p=0.0005 with and without QTL filtering. The positive regulation of amide metabolic process (GO:0034250) was the only GO term that was over-enriched without GWAS filtering, but not with GWAS filtering. There were more overrepresented GO annotations for the height phenotype than for the growth phenotype, possibly because more is known about genes for height. We did not observe significant differences between the annotations enriched in prioritized SNP lists based on QTL filtering and without QTL filtering.

**Table 2.** Experimental summary of executing SSPA on all variants (SNPs) without GWAS filtering based on maximum height phenotype.

| Chromosome Number | Total #SNPs<sup>a</sup> | Prioritized #SNPs<sup>b</sup> | Prioritized #SNPs directly map to known genes | Highest Correlation Coefficient<sup>c</sup> |
| --- | --- | --- | --- | --- |
| 01 | 106,824 | 595 | 259 | 0.64 |
| 02 | 157,148 | 753 | 256 | 0.64 |
| 03 | 132,661 | 656 | 216 | 0.64 |
| 04 | 126,273 | 690 | 208 | 0.64 |
| 05 | 157,930 | 758 | 198 | 0.64 |
| 06 | 157,808 | 649 | 121 | 0.64 |
| 07 | 119,573 | 635 | 136 | 0.64 |
| 08 | 124,899 | 644 | 153 | 0.63 |
| 09 | 124,843 | 679 | 145 | 0.63 |
| 10 | 128,622 | 587 | 171 | 0.63 |
| **"Combined prioritized" SNPs from all chromosomes** | **6646** | **987** | **247** | **0.71** |

<sup>a</sup> Total number of SNPs (corresponding to SNP similarity matrices used as input to the SSPA).
<sup>b</sup> Number of prioritized SNPs extracted using SSPA.
<sup>c</sup> Pearson correlation coefficient of *phenotype similarity matrix* with the *final SNP similarity matrix* (i.e., the summation of SNP similarity matrices corresponding to the prioritized SNPs).

Of the “Top 40” prioritized SNPs (considered a broader range to enhance variance during analysis) extracted with GWAS filtering for both phenotypes at p=0.0001, all except two were in intergenic regions within 700 bp of a known transcript. One SNP, S05_1688138, a deletion event (AATGTG→A) located on Chromosome-5 at 1688138 bp position, was present in the intron of a gene, SORBI_3005G018800, that overlaps with QTLs known to impact dry matter growth rate and plant height (Figure A1). Another SNP, S06_45415791, lies between two pre-miRNA (MIR2118) genes, ENSRNA04948199 and ENSRNA049481970, which are conserved in plants and induce the phased small interfering RNAs (phasiRNAs) production known to impact plant development and fertility (Figure A2; [40]).

A closer look at the 85,709bp region around another intergenic SNP, S01_50208809, shows that it overlaps with QTLs for phenotypes such as plant height, days to flowering, and tiller height. Gene SORBI_3001G265700 flanks this SNP on the right and gene SORBI_3001G265600 on the left. SORBI_3001G265700 encodes a serine-threonine-like kinase and SORBI_3001G265600 encodes an aldolase-1-epimerase like protein (SbA1ELP). The expression profile of the 51 genes that flank and/or overlap these SNPs (according to SnpEff) showed baseline expression of 44 genes in plant structures important for growth and height, such as the root, vegetative meristem, shoot, and stem internodes (Figure B1 (a-c); E-MTAB3839, E-MTAB-4400, and E-GEOD-98817). Included in this list of 51 genes are the genes known to impact transcription factors, cytochrome P450, sugar transporters, protein kinases, Leucine-rich Ser/Thr-kinase membrane proteins, and proteins involved in regulating microtubule binding and assembly that are key to regulating mitosis and growth during the cell cycle and thus height and growth. Most of these genes are expressed across all the experiments (Figure B1(d)).

Of the “Top 10” prioritized SNPs (selected to constrain the analysis to a smaller range) obtained using all variants without GWAS pre-filter, all but three were in intergenic regions within 700 bp of a known transcript. The genes most closely mapping to these prioritized SNPs were different from the genes identified with the GWAS filtering (SORBI_3001G279900, SORBI_3002G119500, SORBI_3005G116400, SORBI_3001G262900, SORBI_3002G153900, SORBI_3002G212200, SORBI_3006G040401, SORBI_3002G132000, SORBI_3007G086800, SORBI_3002G119600, SORBI_3005G116500, SORBI_3007G115500, SORBI_3001G262950, SORBI_3002G154000, SORBI_3006G040500, SORBI_3007G086900).

All these genes overlapped with known QTLs for plant height and most also overlapped with panicle length and panicle emergence QTLs. These panicle traits may also regulate plant height in the reproductive phase. The expression profile of these genes showed baseline expression in plant structures important for growth and height, such as the root, vegetative meristem, shoot, and stem internodes (Figure B2 (a-c); E-MTAB-4400, E-MTAB-3839.and E-GEOD-98817). Two of these genes (SORBI_3001G262950 and SORBI_3006G040401) did not show expression in any of these studies. One gene (SORBI_3005G116400) showed expression in only one study (E-MTAB-4400) where it was expressed in leaf tissues. Most genes are common and expressed in cells of different anatomical parts of the plant and may contribute to overall growth and development (Figure B2(d)).

### 3.5. Comparative Study with Empirical Findings

Further, we validated the prioritized genes obtained for height phenotype as part of our experiment with known height genes in sorghum [41] and a few genes map closely to these (Figure 7). With the GWAS filtering at p-value of 0.0001 with QTL, the only known height gene with a nearby SNP extracted is *Ma1* (SOBIC_006G057866), which has three SNPs nearby and is known to regulate flowering time in *Sorghum* [42] (Figure 7A). It reflects the influence of flowering time on plant height. Without the GWAS filtering, prioritized SNPs are identified near Ma5, Ma2, Ma1, Dw2, and Dw1 (Figure 7B). Recent work on *Sorghum* grown in Ethiopia [43] gives the seven most impactful SNPs on plant height, which fall within known genes (Figure 7A and 7B). With the GWAS filtering at p-value of 0.0001 with QTL, four of these are also near SNPs we identified as prioritized for plant height (SOBIC.003G119600, SOBIC.006G111800, SOBIC.010G085400, and SOBIC.010G066100), two are near known height genes (*DW1*, SOBIC.009G223500 and *Ma5*, SOBIC.001G017500), and one is not near either (SOBIC.006G235400) (Figure 6A). Without the GWAS filtering, an additional SNP identified in the Ethiopian study is also near a SNP identified in this study (Figure 6B).

**Figure 7.**
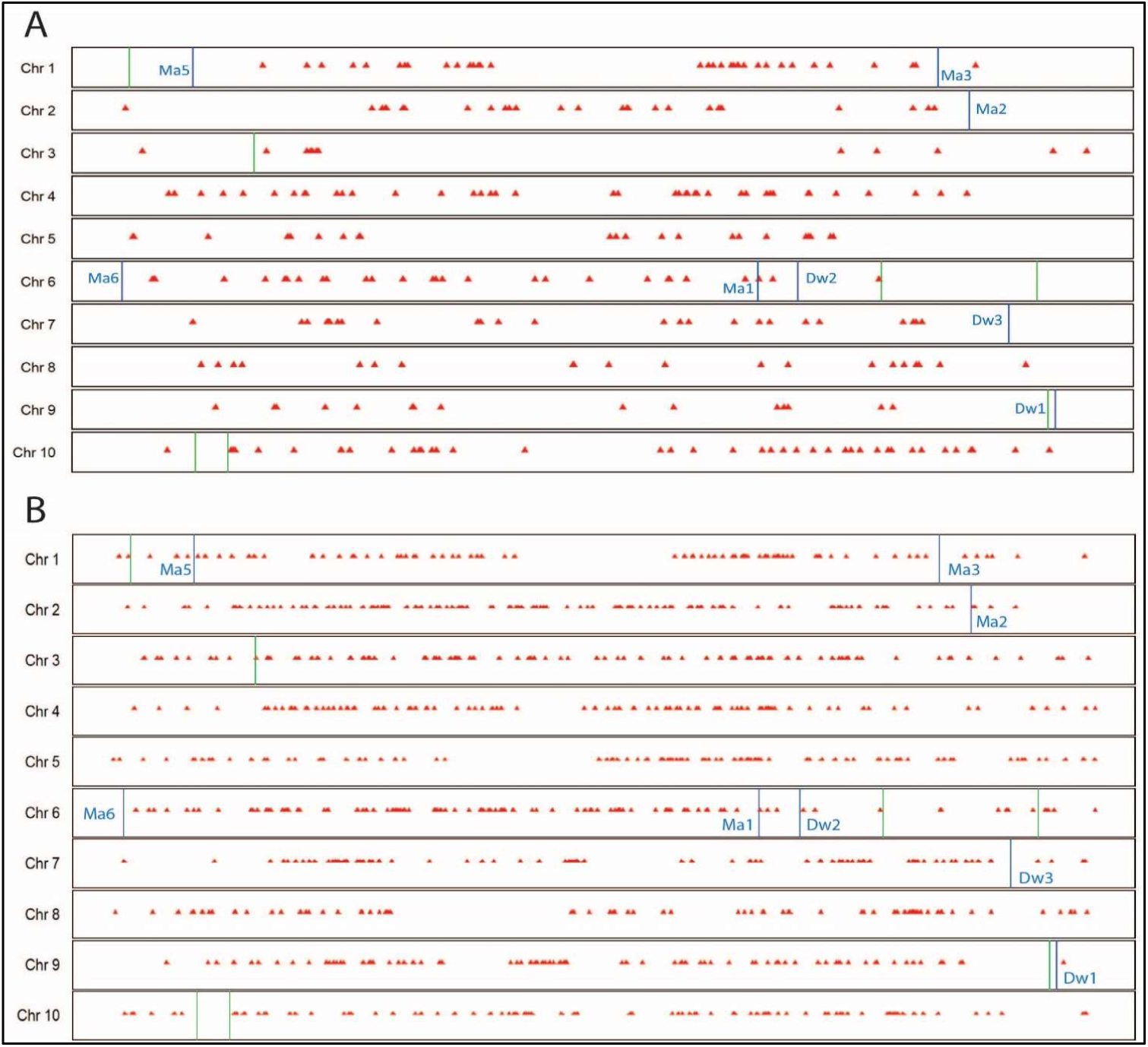
Chromosomal location of prioritized SNPs with known height genes. Prioritized SNPs (red triangles) are scattered throughout the ten chromosomes. The known plant height- associated genes are labeled in blue vertical lines. The green vertical lines show the location of SNPs important for height in sorghum grown in Ethiopia. (A) The 286 SNPs identified for maximum height based on GWAS-filtered set of SNPs with QTL filtering (p-value of .0001) (B) The 987 SNPs identified for maximum height considering all variants without the GWAS filtering.

### 3.6. Generalization Experiment

As described in Section 2.9, we conducted the generalization experiment to evaluate the explanatory power of prioritized SNPs derived from the MAC season 6 dataset, both with and without GWAS filtering (Section 3.1 and 3.2), in the Clemson environment. There were 271 accessions common between the Clemson and MAC Season 6 datasets. The correlation coefficient between Clemson *phenotype similarity matrix* (dimension 271 x 271) and MAC Season 6 *final SNP similarity matrix* (considering 271 instead of 274 accessions) was approximately 0.07 for all experimental conditions with GWAS filtering (Table 1). To compute this, MAC Season 6 *final SNP similarity matrix* was modified by excluding the rows and columns corresponding to the three accessions absent in the Clemson dataset. The correlation coefficient was observed to be 0.06 when considering the *final SNP similarity matrix* obtained from the combined prioritized SNP list without GWAS filtering (Table 2). To further investigate, we examined the histograms of height differences across every pair of accessions for both sites (Figure 8A and 8B), which indicated greater variations in height differences at Clemson compared to MAC Season 6.^4^ Figure 8C highlights the differences in height differences between these two sites, revealing that approximately 55% of the pairs of accessions exhibited differences greater than equal to 50.

**Figure 8.**
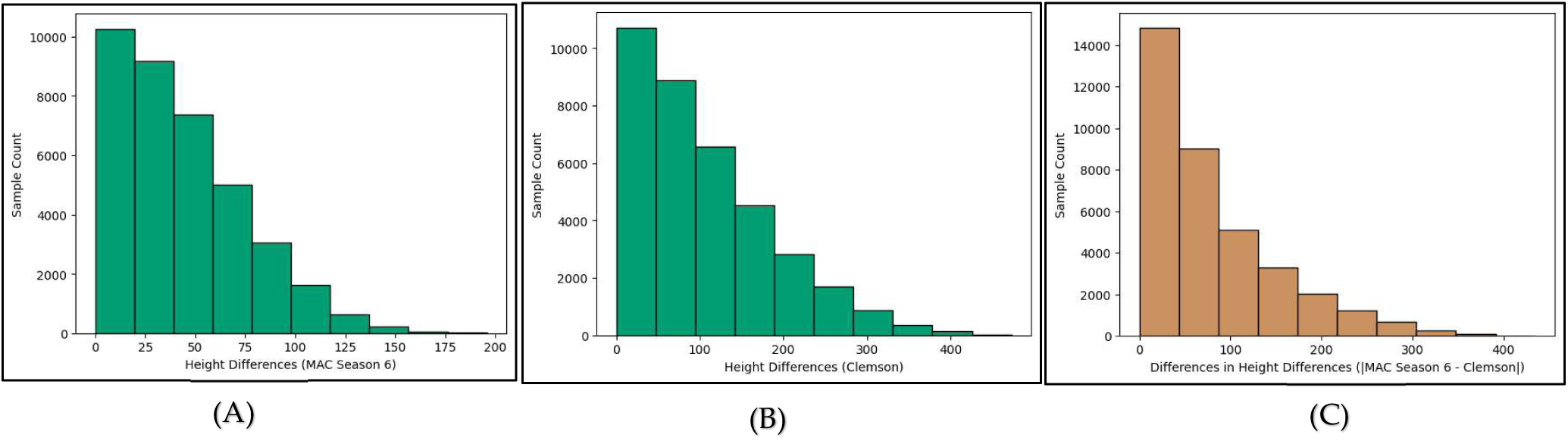
Analysis of differences in height measurements across every pair of accession for MAC Season 6 and Clemson dataset. These diagrams depict the variability in heights in both sites. (A) Height differences in every pair of accessions (274 choose 2) in MAC season 6. (B) Height differences in every pair of accessions (271 choose 2) in Clemson. (C) Difference in height differences between MAC Season 6 and Clemson across every pair of accessions (271 choose 2).

## 4. Discussion

The highest correlation coefficient between *phenotype* and *final SNP similarity matrix* derived from the MAC Season 6 dataset was 0.69 with GWAS filtering (Table 1- Highest Correlation Coefficient) and 0.71 without GWAS pre-filter (Table 2- Highest Correlation Coefficient). These values indicate moderately strong correlations, which are not surprising given the demonstrated correlation of environmental factors on height and growth in other studies [44], because we did not directly capture the effect of the environment in this study. We hypothesize that the total number of SNPs influencing height or growth is much larger than those identified in this study. Depending on the environmental pressures faced by the plant, different sub-groups of SNPs may become predictive. Our results from the Clemson data set (South Carolina) support this. Additionally, the partial overlap in important SNPs with the Ethiopia study may reflect the similarities in hot and arid climates between Ethiopia and Arizona, despite using different accessions. This contrasts with the environmental conditions in South Carolina.

In the absence of empirical data on gene function from sorghum, we can only conclude that this method has the potential to prioritize specific regions of the genome that influence height and growth phenotypes. We found gene homology and expression data supporting the significance of some of the identified sorghum SNPs for the height and growth phenotypes. These SNPs were in the vicinity of gene homologs. For instance, tobacco aldolase-1-epimerase (NbA1ELP) is known to regulate pectin methylesterase (NbPME) antagonistically [45]. Lower expression levels of tobacco NbA1ELP resulted in the dwarf plant phenotype. This suggests that the sorghum ortholog of this tobacco gene (SORBI_3001G265600 SbAiELP) may be associated with plant height and biomass yield in sorghum. Existing GO annotations do not include these traits due to a lack of evidence from sorghum. However, this homologous sorghum gene is expressed in embryo, shoot internode, root, emerging inflorescence, and stem internodes (Figure B1 (a-c)), suggesting that its expression in these tissues is likely to contribute to plant height and growth. Further validation by functional genomics studies is necessary to confirm these findings.

At the outset of this work, we envisaged developing ML models to predict the phenotype based on genomic and environmental information. However, the major difficulty in training ML models with the TERRA-REF dataset was the disparity between the vast size of an entire genome and the limited number of phenotypic observations available. Cultivating on such a large scale to obtain a comparable number of phenotypic measurements with genetic makeup is generally prohibitively expensive, necessitating some form of genomic feature selection. In GWAS, prefiltering (e.g., based on missingness, error probability, linkage disequilibrium, minor allele frequency, or other metrics) can be an effective way to reduce the dimensionality of the datasets (the number of SNPs). However, this approach has shown mixed outcomes [46], and the best method for doing so remains an active area of research.

Our proposed algorithm SSPA can also be utilized to effectively reduce the number of SNPs. This was borne out in our experiment without GWAS filtering that was able to locate several SNPs known to be associated with plant height. These SNPs were not retrieved after applying GWAS with permissive-filtered thresholds that we used in this work. Although the experiment without GWAS filtering does not explicitly account for the population structure, this analysis demonstrated the efficacy of our approach in identifying genes associated with the phenotype of interest. By reducing the number of SNPs, our method facilitates their use as input to the ML models and population structure can be accounted for later by incorporating certain techniques like clustering and principal component analysis in the models [47].

Additionally, we observed that the phenotypic dataset used in our experiment consists of accessions having limited variation in their phenotypic measurements. Consequently, our approach resulted in a phenotype similarity matrix where most of the accessions are highly correlated with each other considering the phenotype under study. Figure 9 displays the histograms of phenotype similarity values across every pair of accessions, calculated with the process described in Section 2.3. We hypothesize that the extraction of prioritized SNPs based on phenotypic measurements by our method would be more pronounced if accessions with greater variations in their phenotypic trait measurements were selected.

**Figure 9.**
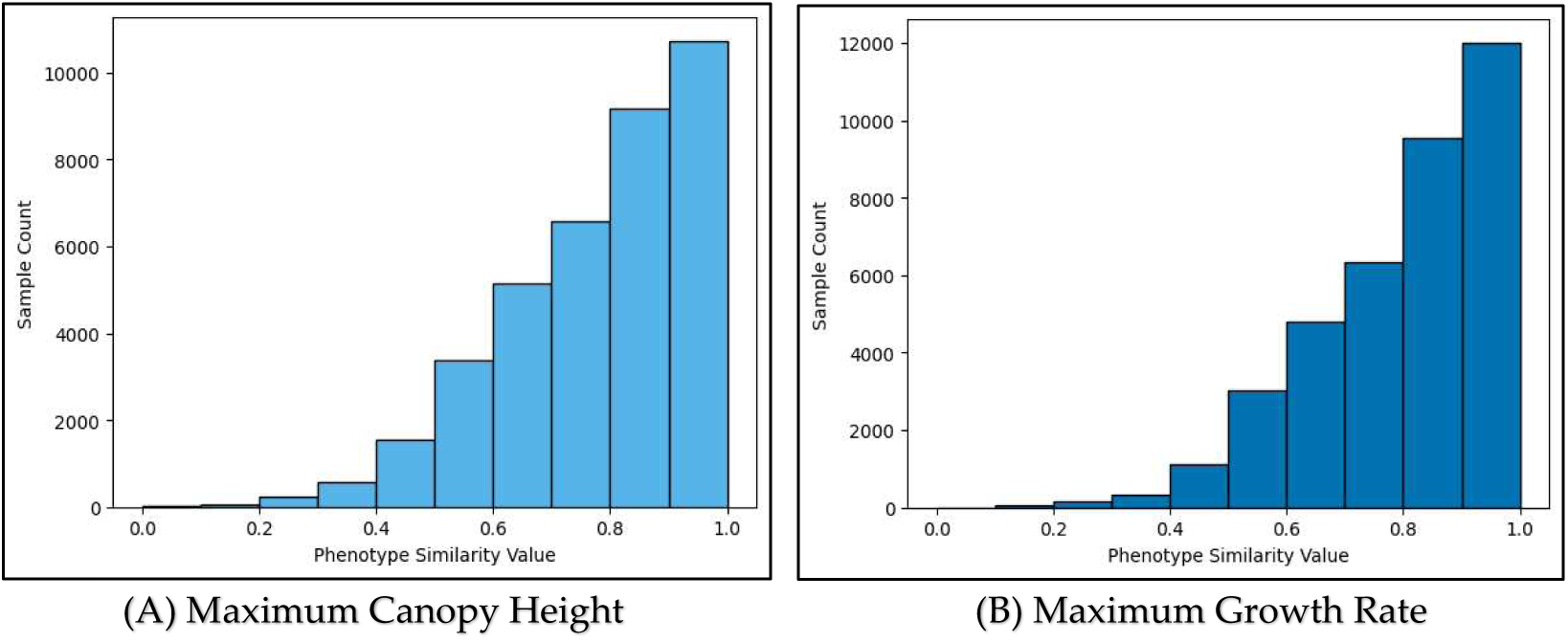
Phenotype similarity value across every pair of accessions in MAC Season 6 dataset. These diagrams depict the estimated phenotype similarity values following the process described in Section 2.3.

## 5. Conclusions

This work focuses on developing an algorithm, named SSPA, for prioritization of SNPs and genes for further analysis to facilitate the domain of functional genetics. Our approach incrementally and cumulatively identifies a set of SNPs that impact a certain phenotypic trait by evaluating both local and global relationships at the pairwise accession level, incorporating a ranking system to prioritize the SNPs. Additionally, this method shows potential for prioritizing genomic regions by reducing the dimensionality (i.e., the number of SNPs) that could be informative for ML/DL methods to predict complex phenotypes. Currently, the method is limited in capturing linear relationships between genotypes and phenotypes, and environmental variations are controlled by normalizing the inputs rather than explicitly incorporating them into the procedure.

In the future, this algorithm could be extended by using weighted correlation metrics to capture non-linear relationships and explicitly incorporating environmental variations into the procedure. Moreover, we used a sequential algorithm for selecting prioritized SNPs, wherein once an SNP is identified as informative, it cannot be removed from the prioritized SNP list. The process of prioritizing SNPs can be further improved by employing other feature selection techniques, such as Sequential Floating Forward Selection (SFFS). We believe that the work reported in this paper provides a novel approach that leverages feature engineering techniques for the prioritization of informative SNPs.

## Author Contributions

Conceptualization, A.R., A.E.T., P.J., D.L. and A.T.; methodology, A.R. and D.P.; validation, A.E.T., P.J. and A.T.; data curation, K.S. and J.G.; writing—original draft preparation, D.P.; writing—review and editing, L.C., P.J., A.E.T. and A.R.; visualization, C.L.; supervision, A.R. and A.E.T.; project administration, L.C.; funding acquisition, A.E.T., A.R., D.L. and P.J. All authors have read and agreed to the published version of the manuscript.

## Data Availability Statement

The original data and code presented in the study are openly available in https://github.com/genophenoenvo/phenotype-to-genotype and in the supplementary files.

## Acknowledgments

We thank the TERRA-REF project for sharing their high throughput phenomics and genomics data.

## Appendix A

**Figure A2.**
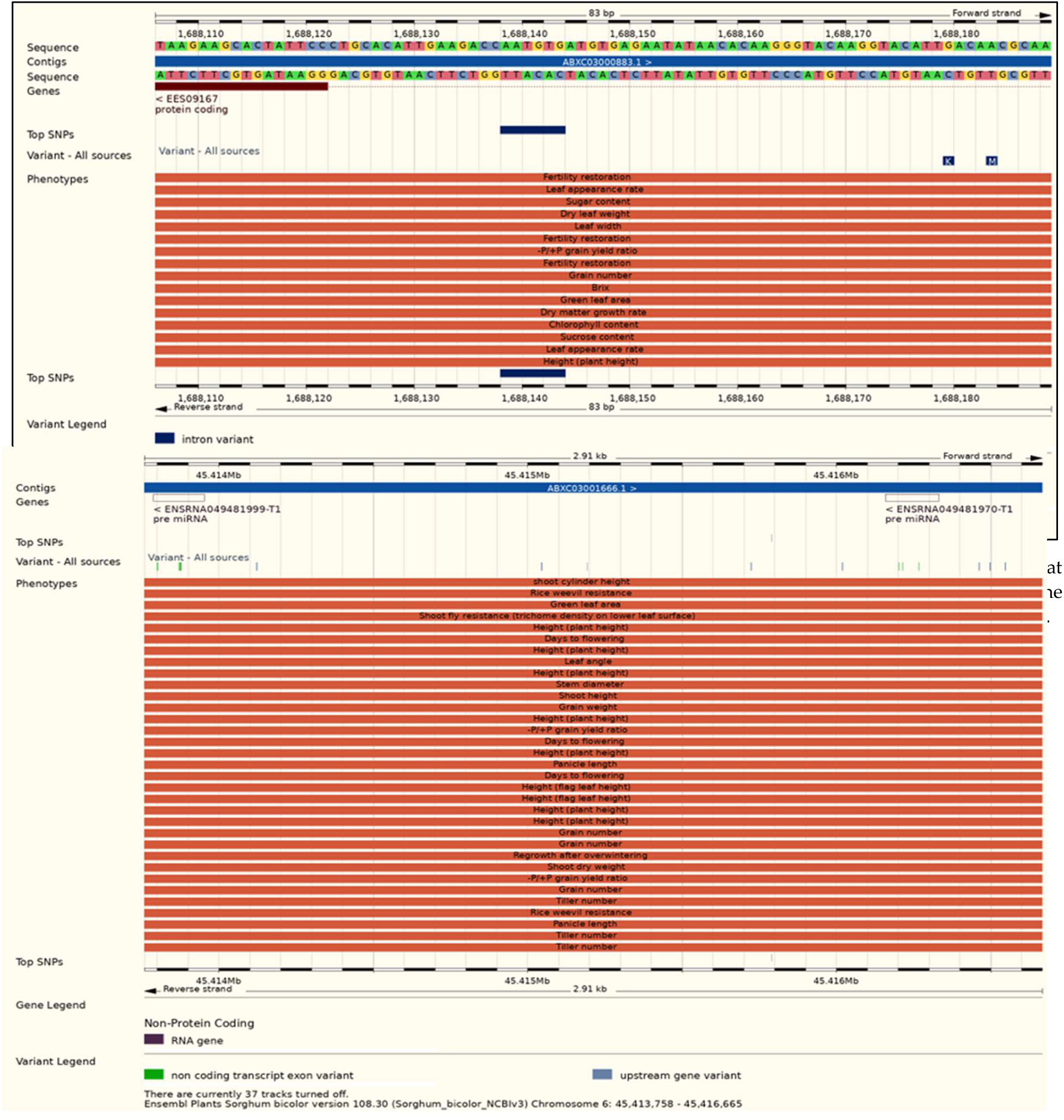
Screenshot from Gramene database. SNP S06_45415791 located on chromosome 6 at 45415791bp position lies between the two pre miRNA (MIR2118) genes ENSRNA049481999 and ENSRNA049481970. miRNA2118 are conserved in plants and induce the phased small interfering RNAs (phasiRNAs) production (Araki et al. 2020). Query performed 15 Dec 2022.

## Appendix B

**Figure B1.**
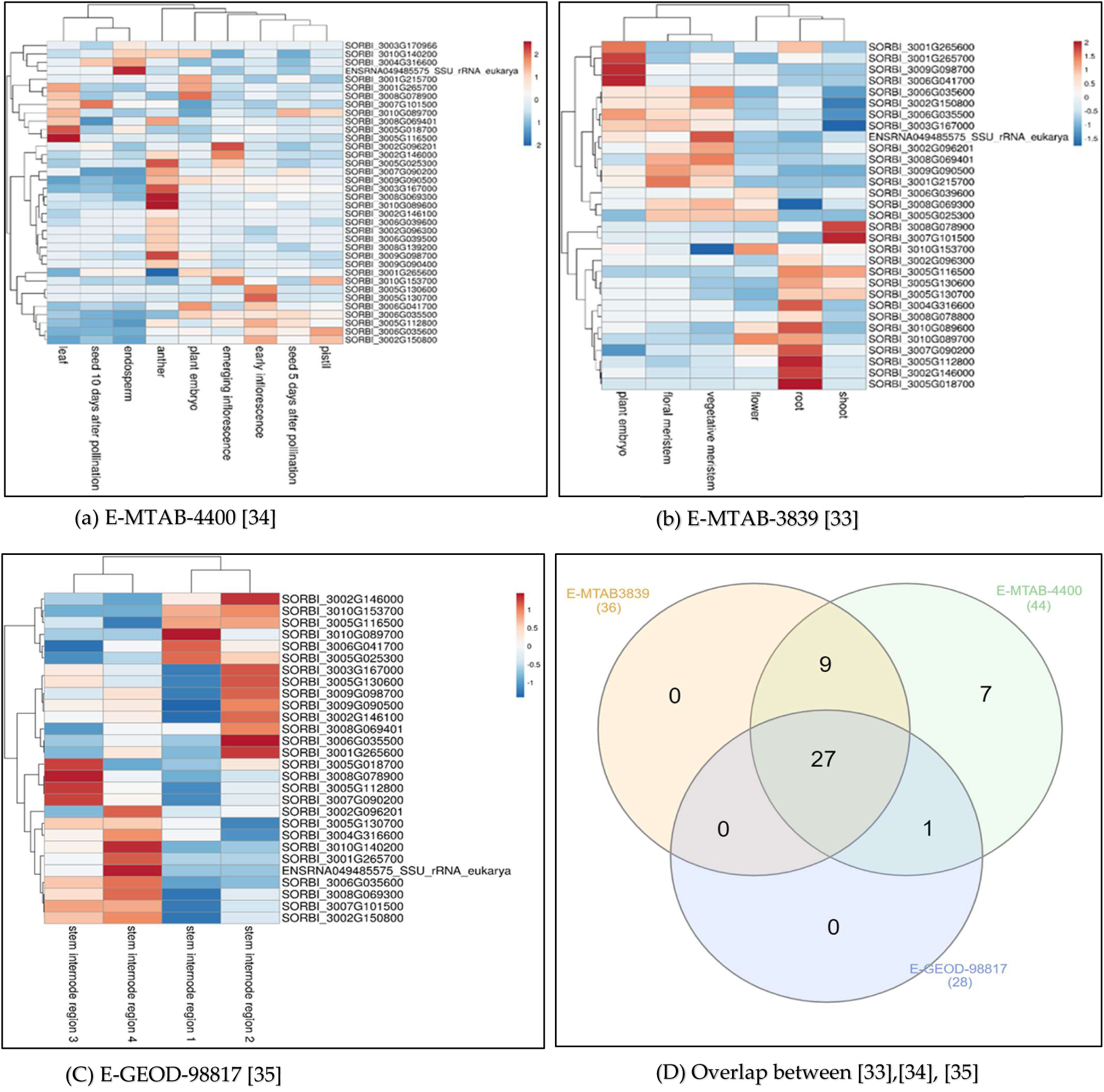
**Gene Expression in Tissues**. We took these 51 genes and queried their baseline expression profile on the EMBL-EBI gene expression atlas. Based on the representative tissue/plant structure atlas studies primarily including vegetative and reproductive growth tissues, we selected three studies to examine using a heatmap visualization, shown in (a), (b), (c). (d) indicates that majority were common genes, suggesting that these genes are expressed in cells of different anatomical parts of the plant and may contribute to overall growth and development.

**Figure B2.**
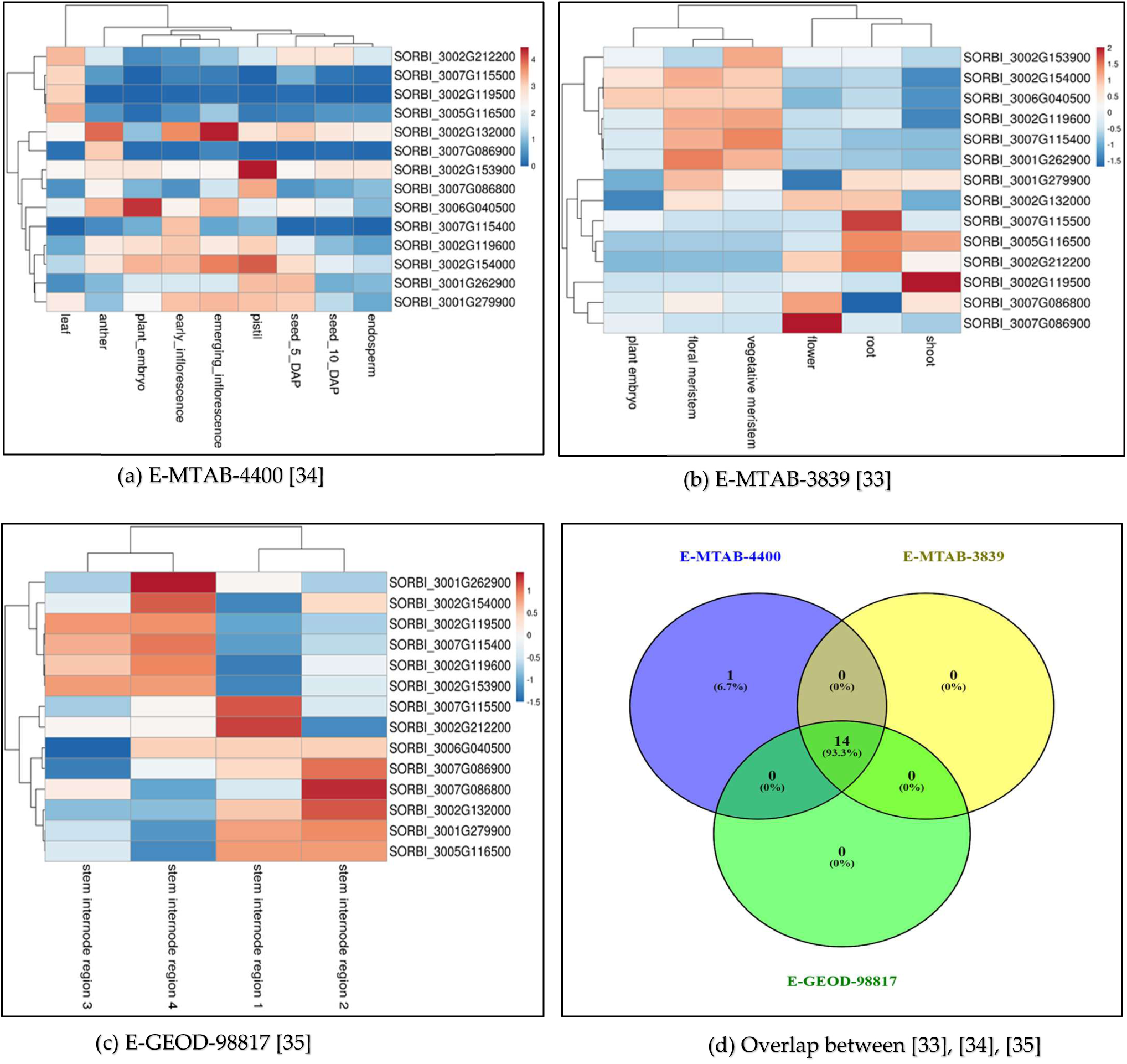
Gene Expression in Tissues. We took these 17 genes and queried their baseline expression profile on the EMBL-EBI gene expression atlas. Based on the representative tissue/plant structure atlas studies primarily including vegetative and reproductive growth tissues, we selected three studies to examine using a heatmap visualization, shown in (a), (b), (c). (d) indicates that majority were common genes, suggesting that these genes are expressed in cells of different anatomical parts of the plant and may contribute to overall growth and development. The one gene in the EMTAB- 4400 was expressed only in the leaf.

## Disclaimer/Publisher’s Note

The statements, opinions and data contained in all publications are solely those of the individual author(s) and contributor(s) and not of MDPI and/or the editor(s). MDPI and/or the editor(s) disclaim responsibility for any injury to people or property resulting from any ideas, methods, instructions or products referred to in the content.

## Footnotes

1 https://github.com/genophenoenvo/phenotype-to-genotype

2 The common set of accessions was considered for future work.

3 For Clemson, we were unable to apply the logistic growth curve model for normalization due to certain environmental factors; however, these measurements are assumed to be comparable with the normalized phenotypic trait measurements of MAC.

4 MAC season 6 height measuresments are normalized through logistic growth curve, whereas for Clemson, end-of-season height measurements were used.

## References

1. Visscher, P.M.; Brown, M.A.; McCarthy, M.I.; Yang, J. Five Years of GWAS Discovery. The American Journal of Human Genetics 2012, 90, 7–24, doi:10.1016/j.ajhg.2011.11.029.

2. Hebbring, S.J. The Challenges, Advantages and Future of Phenome-wide Association Studies. Immunology 2014, 141, 157–165, doi:10.1111/imm.12195.

3. Moore, J.H.; Asselbergs, F.W.; Williams, S.M. Bioinformatics Challenges for Genome-Wide Association Studies. Bioinformatics 2010, 26, 445–455, doi:10.1093/bioinformatics/btp713.

4. Hou, L.; Zhao, H. A Review of Post-GWAS Prioritization Approaches. Front. Genet. 2013, 4, doi:10.3389/fgene.2013.00280.

5. Gupta, P.K.; Kulwal, P.L.; Jaiswal, V. Association Mapping in Plants in the Post-GWAS Genomics Era. In Advances in Genetics; Elsevier, 2019; Vol. 104, pp. 75–154 ISBN 978-0-12-817161-5.

6. Cai, Z.; Guldbrandtsen, B.; Lund, M.S.; Sahana, G. Prioritizing Candidate Genes Post-GWAS Using Multiple Sources of Data for Mastitis Resistance in Dairy Cattle. BMC Genomics 2018, 19, 656, doi:10.1186/s12864-018-5050-x.

7. Marina, H.; Pelayo, R.; Suárez-Vega, A.; Gutiérrez-Gil, B.; Esteban-Blanco, C.; Arranz, J.J. Genome-Wide Association Studies (GWAS) and Post-GWAS Analyses for Technological Traits in Assaf and Churra Dairy Breeds. Journal of Dairy Science 2021, 104, 11850–11866, doi:10.3168/jds.2021-20510.

8. Nicholls, H.L.; John, C.R.; Watson, D.S.; Munroe, P.B.; Barnes, M.R.; Cabrera, C.P. Reaching the End-Game for GWAS: Machine Learning Approaches for the Prioritization of Complex Disease Loci. Front. Genet. 2020, 11, 350, doi:10.3389/fgene.2020.00350.

9. Zhang, Q.; Zhang, Q.; Jensen, J. Association Studies and Genomic Prediction for Genetic Improvements in Agriculture. Front. Plant Sci. 2022, 13, 904230, doi:10.3389/fpls.2022.904230.

10. Burnette, M.; Kooper, R.; Maloney, J.D.; Rohde, G.S.; Terstriep, J.A.; Willis, C.; Fahlgren, N.; Mockler, T.; Newcomb, M.; Sagan, V.;, et al. TERRA-REF Data Processing Infrastructure. In Proceedings of the Proceedings of the Practice and Experience on Advanced Research Computing; ACM: Pittsburgh PA USA, July 22 2018; pp. 1–7.

11. LeBauer, D.; Maxwell, B.; Demieville, J.; Fahlgren, N.; French, A.; Garnett, R.; Hu, Z.; Huynh, K.; Kooper, R.; Li, Z.;, et al. Data From: TERRA-REF, An Open Reference Data Set from High Resolution Genomics, Phenomics, and Imaging Sensors 2020, 1509720474 bytes.

12. Brenton, Z.W.; Cooper, E.A.; Myers, M.T.; Boyles, R.E.; Shakoor, N.; Zielinski, K.J.; Rauh, B.L.; Bridges, W.C.; Morris, G.P.; Kresovich, S. A Genomic Resource for the Development, Improvement, and Exploitation of Sorghum for Bioenergy. Genetics 2016, 204, 21–33, doi:10.1534/genetics.115.183947.

13. Mcmaster, G. Growing Degree-Days: One Equation, Two Interpretations. Agricultural and Forest Meteorology 1997, 87, 291–300, doi:10.1016/S0168-1923(97)00027-0.

14. Plummer, M. JAGS: A Program for Analysis of Bayesian Graphical Models Using Gibbs Sampling. In Proceedings of the Proceedings of the 3rd International Workshop on Distributed Statistical Computing.

15. Plummer, M. Rjags: Bayesian Graphical Models Using MCMC 2008, 4–15.

16. R Core Team (2021). R: A Language and Environment for Statistical Computing. R Foundation for Statistical Computing, Vienna, Austria.

17. Gelman, A.; Rubin, D.B. Inference from Iterative Simulation Using Multiple Sequences. Statist. Sci. 1992, 7, doi:10.1214/ss/1177011136.

18. Guo, J.; LeBauer, D. Genophenoenvo/JAGS-Logistic-Growth: V0.1.0 2022.

19. Goodstein, D.M.; Shu, S.; Howson, R.; Neupane, R.; Hayes, R.D.; Fazo, J.; Mitros, T.; Dirks, W.; Hellsten, U.; Putnam, N.;, et al. Phytozome: A Comparative Platform for Green Plant Genomics. Nucleic Acids Research 2012, 40, D1178–D1186, doi:10.1093/nar/gkr944.

20. Yin, L.; Zhang, H.; Tang, Z.; Xu, J.; Yin, D.; Zhang, Z.; Yuan, X.; Zhu, M.; Zhao, S.; Li, X.;, et al. rMVP: A Memory-Efficient, Visualization-Enhanced, and Parallel-Accelerated Tool for Genome-Wide Association Study. Genomics, Proteomics & Bioinformatics 2021, 19, 619–628, doi:10.1016/j.gpb.2020.10.007.

21. Liu, X.; Huang, M.; Fan, B.; Buckler, E.S.; Zhang, Z. Iterative Usage of Fixed and Random Effect Models for Powerful and Efficient Genome-Wide Association Studies. PLoS Genet 2016, 12, e1005767, doi:10.1371/journal.pgen.1005767.

22. Mace, E.; Innes, D.; Hunt, C.; Wang, X.; Tao, Y.; Baxter, J.; Hassall, M.; Hathorn, A.; Jordan, D. The Sorghum QTL Atlas: A Powerful Tool for Trait Dissection, Comparative Genomics and Crop Improvement. Theor Appl Genet 2019, 132, 751–766, doi:10.1007/s00122-018-3212-5.

23. Pudil, P.; Novovičová, J.; Kittler, J. Floating Search Methods in Feature Selection. Pattern Recognition Letters 1994, 15, 1119–1125, doi:10.1016/0167-8655(94)90127-9.

24. Cingolani, P.; Platts, A.; Wang, L.L.; Coon, M.; Nguyen, T.; Wang, L.; Land, S.J.; Lu, X.; Ruden, D.M. A Program for Annotating and Predicting the Effects of Single Nucleotide Polymorphisms, SnpEff: SNPs in the Genome of Drosophila Melanogaster Strain w ^1118^ ; Iso-2; Iso-3. Fly 2012, 6, 80–92, doi:10.4161/fly.19695.

25. Tello-Ruiz, M.K.; Naithani, S.; Gupta, P.; Olson, A.; Wei, S.; Preece, J.; Jiao, Y.; Wang, B.; Chougule, K.; Garg, P.;, et al. Gramene 2021: Harnessing the Power of Comparative Genomics and Pathways for Plant Research. Nucleic Acids Research 2021, 49, D1452– D1463, doi:10.1093/nar/gkaa979.

26. Cooper, L.; Meier, A.; Laporte, M.-A.; Elser, J.L.; Mungall, C.; Sinn, B.T.; Cavaliere, D.; Carbon, S.; Dunn, N.A.; Smith, B.;, et al. The Planteome Database: An Integrated Resource for Reference Ontologies, Plant Genomics and Phenomics. Nucleic Acids Research 2018, 46, D1168–D1180, doi:10.1093/nar/gkx1152.

27. The Gene Ontology Consortium; Carbon, S.; Douglass, E.; Good, B.M.; Unni, D.R.; Harris, N.L.; Mungall, C.J.; Basu, S.; Chisholm, R.L.; Dodson, R.J.;, et al. The Gene Ontology Resource: Enriching a GOld Mine. Nucleic Acids Research 2021, 49, D325–D334, doi:10.1093/nar/gkaa1113.

28. Thomas, P.D.; Ebert, D.; Muruganujan, A.; Mushayahama, T.; Albou, L.; Mi, H. PANTHER : Making Genome-scale Phylogenetics Accessible to All. Protein Science 2022, 31, 8–22, doi:10.1002/pro.4218.

29. Mi, H.; Muruganujan, A.; Ebert, D.; Huang, X.; Thomas, P.D. PANTHER Version 14: More Genomes, a New PANTHER GO-Slim and Improvements in Enrichment Analysis Tools. Nucleic Acids Research 2019, 47, D419–D426, doi:10.1093/nar/gky1038.

30. Mitchell, A.L.; Attwood, T.K.; Babbitt, P.C.; Blum, M.; Bork, P.; Bridge, A.; Brown, S.D.; Chang, H.-Y.; El-Gebali, S.; Fraser, M.I.;, et al. InterPro in 2019: Improving Coverage, Classification and Access to Protein Sequence Annotations. Nucleic Acids Research 2019, 47, D351–D360, doi:10.1093/nar/gky1100.

31. Tello-Ruiz, M.K.; Naithani, S.; Stein, J.C.; Gupta, P.; Campbell, M.; Olson, A.; Wei, S.; Preece, J.; Geniza, M.J.; Jiao, Y.;, et al. Gramene 2018: Unifying Comparative Genomics and Pathway Resources for Plant Research. Nucleic Acids Research 2018, 46, D1181–D1189, doi:10.1093/nar/gkx1111.

32. Papatheodorou, I.; Fonseca, N.A.; Keays, M.; Tang, Y.A.; Barrera, E.; Bazant, W.; Burke, M.; Füllgrabe, A.; Fuentes, A.M.-P.; George, N.;, et al. Expression Atlas: Gene and Protein Expression across Multiple Studies and Organisms. Nucleic Acids Research 2018, 46, D246–D251, doi:10.1093/nar/gkx1158.

33. Olson, A.; Klein, R.R.; Dugas, D.V.; Lu, Z.; Regulski, M.; Klein, P.E.; Ware, D. Expanding and Vetting *Sorghum Bicolor* Gene Annotations through Transcriptome and Methylome Sequencing. The Plant Genome 2014, 7, plantgenome2013.08.0025, doi:10.3835/plantgenome2013.08.0025.

34. Davidson, R.M.; Gowda, M.; Moghe, G.; Lin, H.; Vaillancourt, B.; Shiu, S.; Jiang, N.; Robin Buell, C. Comparative Transcriptomics of Three Poaceae Species Reveals Patterns of Gene Expression Evolution. The Plant Journal 2012, 71, 492–502, doi:10.1111/j.1365-313X.2012.05005.x.

35. Kebrom, T.H.; McKinley, B.; Mullet, J.E. Dynamics of Gene Expression during Development and Expansion of Vegetative Stem Internodes of Bioenergy Sorghum. Biotechnol Biofuels 2017, 10, 159, doi:10.1186/s13068-017-0848-3.

36. Metsalu, T.; Vilo, J. ClustVis: A Web Tool for Visualizing Clustering of Multivariate Data Using Principal Component Analysis and Heatmap. Nucleic Acids Res 2015, 43, W566–W570, doi:10.1093/nar/gkv468.

37. McLaren, W.; Gil, L.; Hunt, S.E.; Riat, H.S.; Ritchie, G.R.S.; Thormann, A.; Flicek, P.; Cunningham, F. The Ensembl Variant Effect Predictor. Genome Biol 2016, 17, 122, doi:10.1186/s13059-016-0974-4.

38. Naithani, S.; Geniza, M.; Jaiswal, P. Variant Effect Prediction Analysis Using Resources Available at Gramene Database. In Plant Genomics Databases; Van Dijk, A.D.J., Ed.; Methods in Molecular Biology; Springer New York: New York, NY, 2017; Vol. 1533, pp. 279–297 ISBN 978-1-4939-6656-1.

39. Lex, A.; Gehlenborg, N.; Strobelt, H.; Vuillemot, R.; Pfister, H. UpSet: Visualization of Intersecting Sets. IEEE Trans. Visual. Comput. Graphics 2014, 20, 1983–1992, doi:10.1109/TVCG.2014.2346248.

40. Araki, S.; Le, N.T.; Koizumi, K.; Villar-Briones, A.; Nonomura, K.-I.; Endo, M.; Inoue, H.; Saze, H.; Komiya, R. miR2118- Dependent U-Rich phasiRNA Production in Rice Anther Wall Development. Nat Commun 2020, *11*, 3115, doi:10.1038/s41467-020-16637-3.

41. Quinby, J.R.; Karper, R.E. Inheritance of Height in Sorghum ^1^. Agronomy Journal 1954, 46, 211–216, doi:10.2134/agronj1954.00021962004600050007x.

42. Quinby, J.R.; Karper, R.E. The Inheritance of Three Genes That Influence Time of Floral Initiation and Maturity Date in Milo ^1^. Agronomy Journal 1945, 37, 916–936, doi:10.2134/agronj1945.00021962003700110006x.

43. Enyew, M.; Feyissa, T.; Carlsson, A.S.; Tesfaye, K.; Hammenhag, C.; Seyoum, A.; Geleta, M. Genome-Wide Analyses Using Multi-Locus Models Revealed Marker-Trait Associations for Major Agronomic Traits in Sorghum Bicolor. Front. Plant Sci. 2022, 13, 999692, doi:10.3389/fpls.2022.999692.

44. Tari, I.; Laskay, G.; Takács, Z.; Poór, P. Response of Sorghum to Abiotic Stresses: A Review. J Agronomy Crop Science 2013, 199, 264–274, doi:10.1111/jac.12017.

45. Sheshukova, E.V.; Komarova, T.V.; Pozdyshev, D.V.; Ershova, N.M.; Shindyapina, A.V.; Tashlitsky, V.N.; Sheval, E.V.; Dorokhov, Y.L. The Intergenic Interplay between Aldose 1-Epimerase-Like Protein and Pectin Methylesterase in Abiotic and Biotic Stress Control. Front. Plant Sci. 2017, 8, 1646, doi:10.3389/fpls.2017.01646.

46. Danilevicz, M.F.; Gill, M.; Anderson, R.; Batley, J.; Bennamoun, M.; Bayer, P.E.; Edwards, D. Plant Genotype to Phenotype Prediction Using Machine Learning. Front. Genet. 2022, 13, 822173, doi:10.3389/fgene.2022.822173.

47. López-Cortés, X.A.; Matamala, F.; Maldonado, C.; Mora-Poblete, F.; Scapim, C.A. A Deep Learning Approach to Population Structure Inference in Inbred Lines of Maize. Front. Genet. 2020, 11, 543459, doi:10.3389/fgene.2020.543459.

