## Supplementary File for "Post-GWAS Prioritization of Genome-Phenome Association in Sorghum"

### *Supplementary Material*

#### 1 Supplementary Data

- VCF files obtained after GWAS analysis performed using rMVP with the FarmCPU algorithm at various p-value levels and the SNP similarity matrices is available here: https://github.com/genophenoenvo/sorghum_data/releases/tag/v0.0.4
- Chromosome-wise VCF files considering all variants and corresponding SNP similarity matrices is available here: https://storage.googleapis.com/gpe-sorghum/whole-vcf-snp-arrays

#### 2 Supplementary Figures

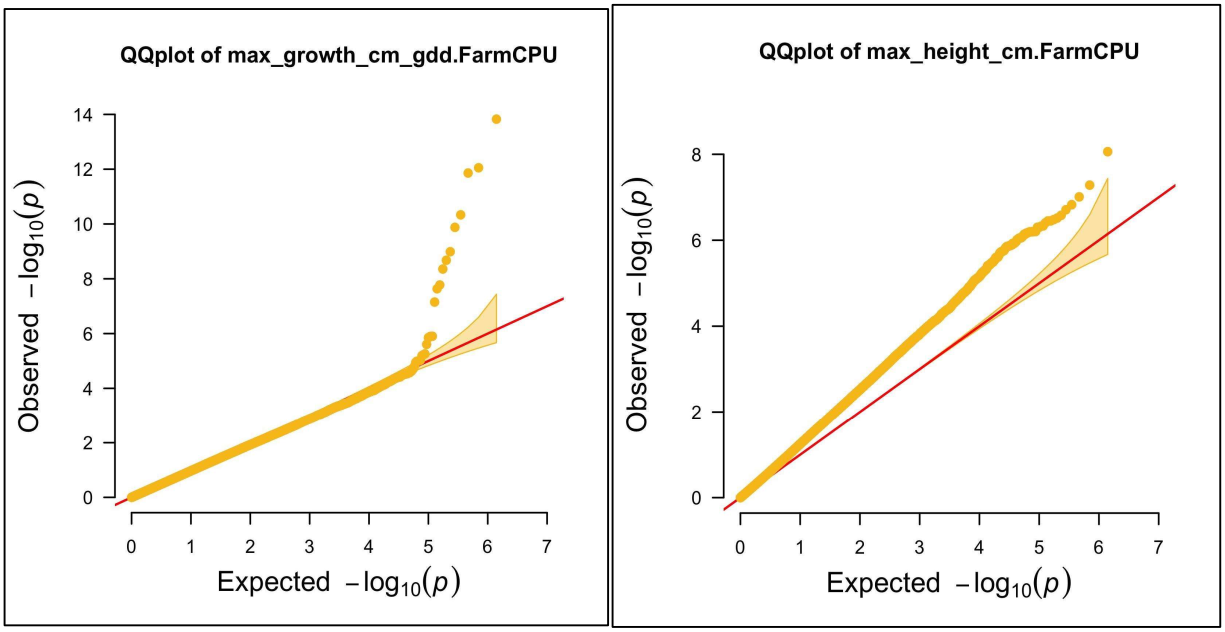


**Supplementary Figure 1.** QQ Plot for the maximum growth rate and maximum height phenotypes in *Sorghum bicolor* "Season 6" TERRA-REF data set.
